# Human Proteome-wide Mechanistic Interpretation of Missense Variants through Protein Feature Enrichment Score

**DOI:** 10.64898/2026.05.26.726248

**Authors:** Seulki Kwon, Jordan Safer, Marina DiStefano, Matthew Lebo, Heidi L. Rehm, Sumaiya Iqbal

## Abstract

Missense variant interpretation remains a central challenge in clinical and medical genetics, with most observed variants being variants of uncertain significance (VUS). Computational variant effect predictors can achieve high pathogenicity classification performance, but without revealing the underlying mechanism and a translatable interpretation. Here we present the Protein Feature Enrichment Score (PFES), which quantifies the molecular context of missense variants through statistical enrichment of 103 protein structural, functional, and physicochemical features across 85,321 pathogenic and 130,719 control variants spanning 20 protein functional classes. We show that the protein feature (PF) enrichment patterns of variants are conserved within functional classes and vary substantially across classes, both in magnitude and directions depending on functional context. PFES not only partitions variants into *PF-Enriched* (pathogenic-like)*, PF-Neutral,* and *PF-Depleted* (benign-like) categories but also provides a mechanistic interpretation by decomposing the score into subscores from biologically interpretable protein feature attributes. We demonstrate that PFES shows a high concordance with VUS reclassification and prioritization: across 596 genes, pathogenicity-leaning VUS-high variants were seven-fold enriched in PF-Enriched variants. PFES decomposition further revealed that loss-of-function and gain-of-function variants are distinguished by disproportionate enrichment of protein-protein interaction features in the latter. We computed PFES across 223 million possible missense variants (17.7% PF-Enriched) and built a publicly available resource that addresses not just whether a variant is pathogenic, but which protein characteristics are disrupted. Proteome-wide application across 20,153 genes prioritizes established rare disease genes and nominates therapeutically amenable targets whose pathogenic variation is driven by interpretable structural and functional protein feature disruption.

**One Sentence Summary:** PFES is a proteome-wide resource to quantify the protein context of missense variants, enabling mechanistically transparent variant interpretation.

## Introduction

Accurately predicting the effects of genetic variants has been a key challenge in genetics and protein biology. Missense variants, where a single amino acid is altered, are the most frequent protein-coding variants linked to numerous diseases and conditions. However, the functional and molecular impact of many of these variants is still unknown; clinically, merely 7% of missense variants are classified as pathogenic or benign, leaving the vast majority as variants of uncertain significance (VUS) (*1*, *2*). The interpretation of DNA variants is also crucial in functional genomics, where understanding how synthetic variations impact biological pathways and cellular phenotypes can reveal insights into disease mechanisms (*3–5*). Tools like deep mutational scanning (*6*, *7*), CRISPR-based base-editing (*8*, *9*), and prime-editing technologies (*10*) now enable the generation of synthetic variants, allowing researchers to systematically study the effects of specific mutations in model systems (*11–13*). These experimental approaches are effective for determining variant effects on a particular phenotype; however, they are system-specific and costly to keep pace with the growing number of clinical variants and the demand for their mechanistic interpretation.

Interpreting the pathogenicity of genetic variants is a complex and challenging task due to the intricate interplay between deleteriousness, conservation, and molecular mechanisms. A variant’s potential to disrupt function and lead to pathogenesis cannot be fully understood by examining these factors in isolation, as they are highly interdependent. Deleteriousness is context-dependent, as a variant in a highly conserved region is more likely to be harmful, yet even in less conserved regions, subtle changes can profoundly affect protein dynamics, allosteric regulation, or interactions with other biomolecules (*14, 15*). Similarly, while conservation indicates evolutionary importance, not all conserved residues contribute equally to protein function, making it difficult to predict which changes are truly pathogenic (*16*). Furthermore, the impact of a protein-coding variant depends on its molecular context, such as its 3D structure, involvement in intra- and inter-protein interactions, and potential to undergo post-translational modifications (PTMs), adding another layer of complexity (*17–19*). As a result, accurate pathogenicity prediction requires a multi-dimensional approach that integrates structural, biochemical, and evolutionary data to capture the nuanced ways in which variants influence protein behavior and ultimately contribute to disease phenotypes.

Current approaches to variant interpretation represent two complementary paradigms, each with distinct strengths and limitations. Computational variant effect predictors (VEP) leverage evolutionary conservation (SIFT (*20*), PROVEAN (*21*)), population frequency (popEVE (*22*), PRESCOTT (*23*)), clinically annotated variants (Polyphen-2 (*24*), VARITY (*25*)), protein structure (SaProt (*26*), SNPs&GO3D(*27*)), and meta-prediction strategies (CADD (*28*), REVEL (*29*)) to provide proteome-wide pathogenicity predictions. Most VEPs are built on machine-learning frameworks, ranging from classical classifiers to deep neural networks and, more recently, protein language models such as AlphaMissense (*30*) or ESM-based methods (*31*), which achieve high prediction accuracy. However, these methods largely function as black boxes, providing predictions without revealing the underlying molecular mechanisms. Conversely, multiplexed assays of variant effect (MAVE) (*1, 32*) provide functional readouts, such as protein activity, abundance, and cellular growth rate, offering protein- and assay-specific mechanistic insights. While MAVEs provide functional measurement, they are limited to specific proteins and assays, and the connection between experimental phenotypes and disease mechanisms can vary.

To achieve mechanistic interpretability at scale, a fundamental question must be addressed: which molecular context contributes to the pathogenicity of a missense variant? Decades of structural biology, biochemistry, and proteomics have produced an extraordinarily rich body of protein annotations, spanning catalytic residues, disordered regions, PTMs, and domain architectures (*33, 34*), defining the sequence-level context at a variant level. Additionally, proteins’ 3D structures are now available at near-proteome scale, powered by recent advances in computational structure prediction, most notably AlphaFold (*35, 36*), alongside experimental structures deposited in the Protein Data Bank (PDB) (*37*). These resources collectively capture the structural, interaction, and functional context of mutations. Yet this wealth of annotations defining the molecular context (sequence-level, structural, and functional) of a variant position has not been integratively leveraged for proteome-wide variant interpretation.

In this study, we address this gap through developing a protein feature enrichment score (PFES), which is an interpretable statistical framework that prioritizes protein sequence, structural, and functional annotations of a variant position towards its mechanistic interpretation. Our approach is grounded in the principle that the multi-dimensional molecular context of a variant determines its consequence: pathogenicity emerges from the intersection of what is being changed, where the change occurs within the protein’s structural architecture, and how that location relates to functional constraints of the protein’s family. A key strength of our framework lies in its systematic integration of comprehensive protein knowledge bases: we compute 3D structural features from experimental and predicted structures (PDB and AlphaFold2), and incorporate extensive sequence and functional annotations from UniProtKB (*34*), PhosphoSitePlus (*33*), and PANTHER (*38*). By quantifying enrichment patterns of diverse protein features of pathogenic versus benign variants across 20 protein classes conserved through evolution for specific functions, our protein feature attribution approach provides mechanistic hypotheses about why a protein variant is disruptive, offering interpretable assessments that complement existing variant effect predictors and assay readouts.

## Results

### Overview of data and Protein Feature Enrichment Score (PFES) framework

To enable mechanistic interpretation of missense variants across the entire human proteome, we developed a framework that quantifies the molecular context of pathogenic and benign variants through the statistical enrichment of protein features. The outline of the bioinformatic workflow is shown in **Fig. 1**. First, we collected missense variants annotated on canonical protein isoforms from ClinVar (*39*), Human Gene Mutation Database (HGMD) (*40*), and Genome Aggregation Database (gnomAD) (*41*) (**Fig. 1a**; **Methods:** Missense variant collection from ClinVar, HGMD, and gnomAD), and mapped them onto each protein’s 3D structure – 85,321 pathogenic variants (from ClinVar pathogenic/likely pathogenic; P/LP) and disease mutations (from HGMD); 104,848 benign variants (from ClinVar benign/likely benign; B/LB), providing crucial negative controls for distinguishing pathogenic molecular signatures; 12,079,818 population variants (from gnomAD), a sample of naturally occurring variants expected to be neutral and not disease-causing, and further divided by their allele frequency (AF) categories: Rare (AF < 0.1%), Intermediate (0.1% ≤ AF < 1%), and Common (AF ≥ 1%); and 1,489,214 variants of uncertain significance (VUS) (from ClinVar) whose clinical interpretation is not complete due to lack of evidence.

**Fig. 1.**
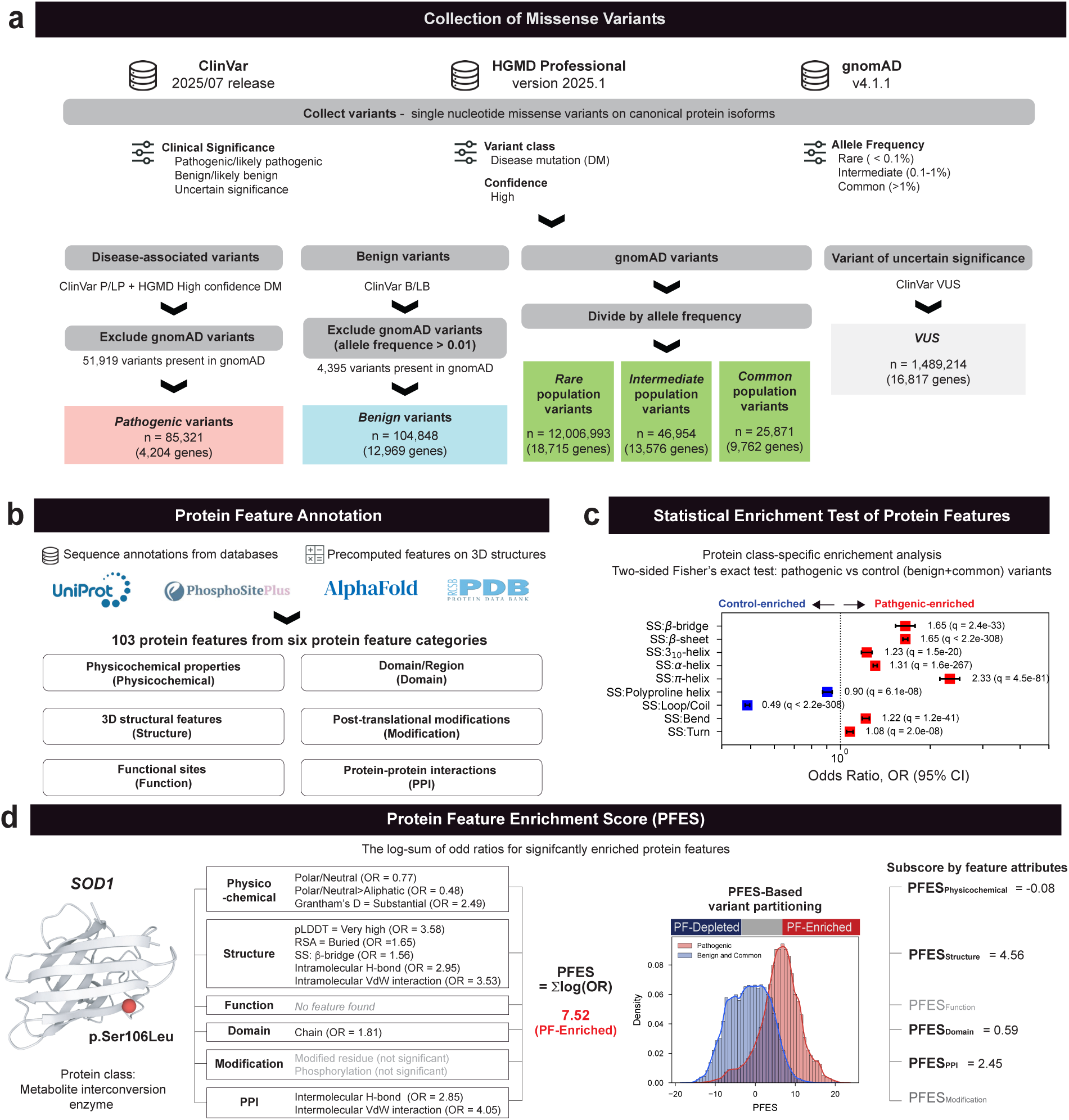
Overview of the PFES framework and bioinformatic workflow. **(a)** Missense variants annotated on canonical protein isoforms were collected from ClinVar (*39*) (2025/07 release), HGMD Professional (*40*) (version 2025.1), and gnomAD (*41*) (v4.1.1) (**Methods**: Missense variant collection from ClinVar, HGMD, and gnomAD). Variants categorized into four groups – pathogenic, benign, population, and VUS – are shown along with the number of variants (n) and corresponding gene counts. All variants were mapped onto protein 3D structures (AlphaFoldDB (*35*) and Protein Data Bank (*37*)). **(b)** Each variant was annotated with protein features spanning six categories: amino acid’s physicochemical properties, 3D structural features, functional site annotations, domain and region annotations, post-translational modification sites, and protein-protein interaction residues. Protein features annotated on sequences were collected from UniProtKB (*34*) and PhosphoSitePlus (*33*) **(c)** Statistical enrichment of protein features were performed by two-sided Fisher’s exact tests. Odds ratios with 95% confidence intervals (CI) are shown for secondary structure features as a representative example, with q-values from FDR correction. Features with OR > 1 are enriched among pathogenic variants (red); features with OR < 1 are enriched among controls (blue). **(d)** (*Left*) PFES computation illustrated for *SOD1* p.Ser106Leu, showing annotated protein features and their respective odds ratios. Significantly enriched features from each of the six feature categories contribute their log-odds ratios to the total PFES (= 7.52). (*Right*) Empirical distributions of pathogenic and control datasets, which are used to partition variants by PFES into PF Depleted, Neutral, and Enriched variants, are shown. Based on the empirical PFES distributions, p.Ser106Leu is classified as PF-Enriched. Subscores decomposed by feature attributes are shown on the right.

Next, each variant was annotated with 103 protein features capturing six aspects of the molecular context: physicochemical properties of amino acids, 3D structural features, domain/region annotations, functional sites (i.e., active sites, binding sites, DNA-binding sites), post-translational modification sites, and protein-protein interaction (PPI) residues, collected from public databases or precomputed (**Fig. 1b**, **Methods**: Protein feature annotations). Comparison of these protein feature distributions across pathogenic, benign, and population, described in the following section, informed the selection of the control dataset (benign and common population variants) against the case (pathogenic) dataset for a statistical enrichment test to maximize the distinction between disease-associated and neutral variants. Statistical enrichment tests were then performed on this finalized dataset to characterize statistically significant protein features enriched in case versus control variants (**Methods**: Fisher’s exact test and Benjamini-Hochberg procedure). **Fig. 1c** shows a representative example using secondary structure features, where odds ratios (OR) from two-sided Fisher’s exact tests reveal differential enrichment between the given case (i.e., pathogenic) and control (i.e., benign and common population) variants – structured elements such as β-sheet (OR=1.65, corrected p, q < 0.05) or α-helices (OR=1.31, q < 0.05) are enriched among pathogenic variants, while loop/coiled regions are enriched among controls, illustrating how individual protein features carry directional signals.

From the resulting odds ratios, we defined a protein feature enrichment score (PFES) as the sum of log-odds ratios of protein features showing significant enrichment among pathogenic or benign datasets, providing a cumulative measure of molecular context for a given missense variant (**Methods**: Computation of Protein Feature Enrichment Score (PFES)). The score enables two key utilities for missense variant interpretation. First, by comparing a variant’s PFES against empirical distributions of pathogenic and benign variants, variants are partitioned into three categories: PF-Enriched, in which the variant’s protein features (PF) resemble pathogenic variants and is statistically distinct from benign variants; PF-Neutral, in which the molecular profile is consistent with both pathogenic and benign distributions, lacking a distinctive signal in either direction; or PF-Depleted, in which the molecular profile resembles benign variants and is statistically distinct from pathogenic variants (**Methods**: PFES-based partitioning of PF-Enriched, Neutral, and Depleted variants). Second, PFES decomposes into subscores from six different protein feature groups (score attributes), revealing which protein features drive the partitioning and providing a mechanistic basis for interpretation. For example, **Fig. 1d** (*left*) illustrates the computation of the PFES of SOD1:p.Ser106Leu that has significantly enriched features by summing their log-odds ratios, resulting in a score of 7.52. Based on the empirical distribution of PFES, we categorized it as PF-Enriched, indicating that this variant’s protein characteristics resemble known pathogenic variants and are statistically distinct from benign variants. Score decomposition (**Fig. 1d**, *right*) further reveals that structural features and protein-protein interaction (PPI) are the primary contributors (PFES_Structure_ = 4.56 and PFES_PPI_ = 2.45). Individual score attributes can similarly be partitioned into PF-Enriched/Neutral/Depleted based on their own empirical distributions, which will be discussed in later sections. Together, empirical partitioning and mechanistic decomposition distinguish PFES from conventional pathogenicity scores that output a single prediction without molecular explanation. For the rest of the paper, we further describe the PFES framework, use it to characterize protein feature enrichment across protein functional classes, and then demonstrate its utility for mechanistic interpretation of variants, benchmarking against functional assays, and prioritizing variants of uncertain significance (VUS).

### Common population variants as proxy-benign controls from comparative protein feature analysis

Pathogenic and benign variants are known to differ in their effects on protein features (*18, 42, 43*).In addition to clinically identified benign variants, population variants observed in relatively healthy individuals through large-scale sequencing datasets, such as gnomAD, are often used as proxy-benign data in comparative studies (*44–46*). Their appearance in the general population suggests they are largely tolerated. However, variants across the allele frequency (AF) spectrum may exhibit different protein feature profiles.

To test this, we performed a comparative analysis of the distribution of protein features between pathogenic (from ClinVar and HGMD), benign (from ClinVar), and population variants across different allele frequency groups from gnomAD (rare, intermediate, and common). We quantified pairwise differences using the Kolmogorov-Smirnov (KS) statistic for continuous features, such as AlphaFold pLDDT, and Cohen’s w for binary and categorical features, such as domain annotation or secondary structures (**Methods:** Pairwise feature distribution comparison: Kolmogorov-Smirnov test and Cohen’s w). Both metrics range from 0 (identical distributions) to 1 (maximal divergence), providing a unified scale for comparing distributional differences across feature types.

**Fig. 2a** shows the distributions of relative surface area (RSA) across five different datasets. Pathogenic variants exhibited a sharp peak at near-zero RSA values, indicating they are predominantly buried in the protein core. In contrast, benign variants exhibited a broader distribution with higher density around moderate RSA values (0.4-0.8), suggesting they commonly occur in solvent-exposed regions. Among population variants, rare variants show a weakly bimodal pattern, with a peak at low RSA overlapping pathogenic variants and a broader peak at high RSA aligning with benign variants, while common variants shift toward higher RSA values, aligning with benign variants. Pairwise KS statistics (**Fig. 2a**, *right*) quantify those differences: pathogenic variants were the most distinct from all other groups (KS > 0.3), while ClinVar benign variants showed the smallest divergence with common population variants (KS = 0.036). Similar analyses using a binary feature such as intramolecular van der Waals (VdW) interactions showed the same trend: common population variants were compositionally most like clinically annotated benign variants (Cohen’s w = 0.113; **Fig. 2b**). **Fig. 2c** shows that this pattern holds across all features in general, where benign variants exhibited the smallest mean KS statistic (continuous features, *left*) and mean Cohen’s w (categorical features, *right*) with common population variants compared to others (**Supplementary Fig. 1**).

**Fig. 2.**
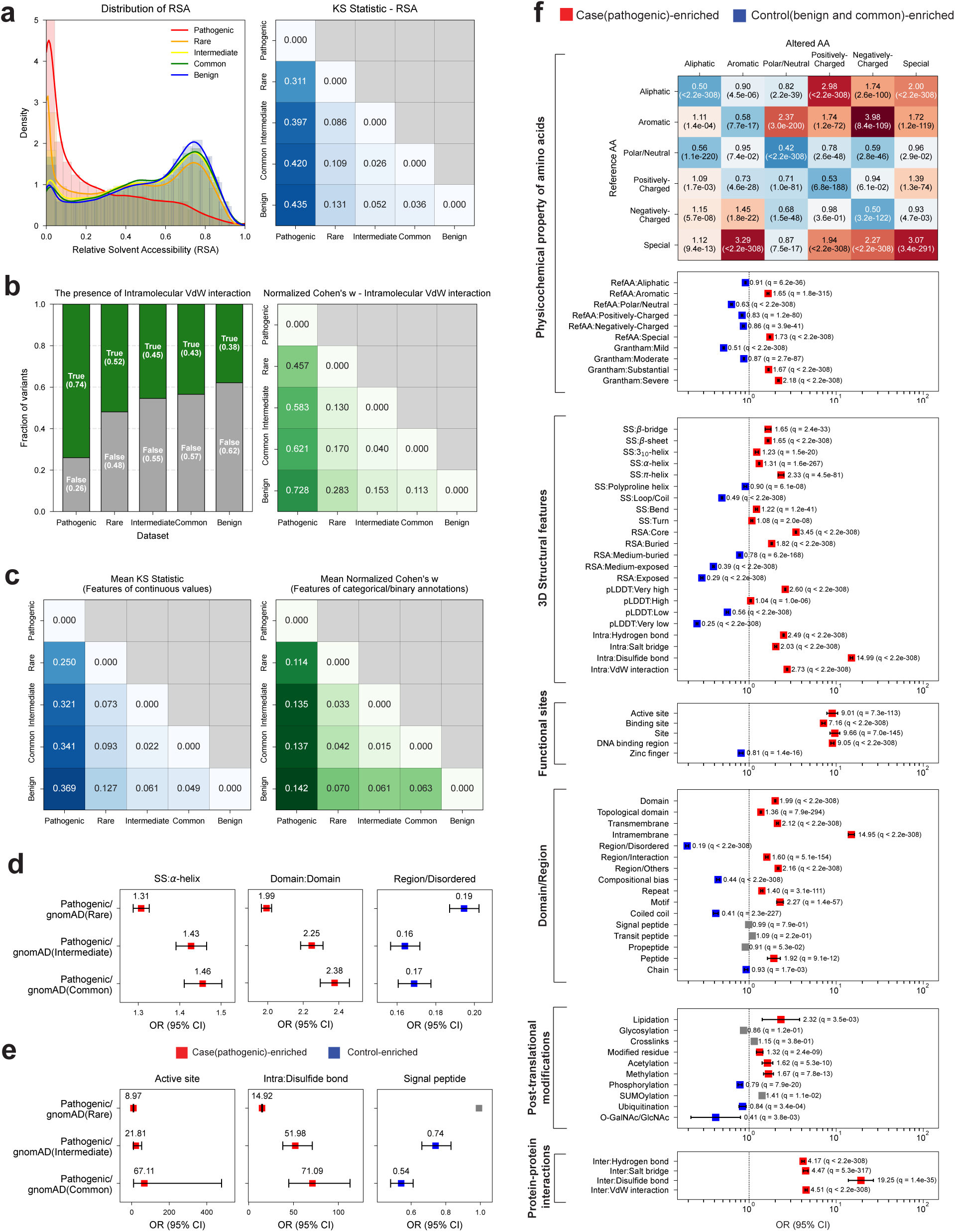
Comparative analysis of protein feature distributions across variant datasets informs control group selection. (a) (*Left*) Density distributions of relative solvent accessibility (RSA) across five variant datasets: pathogenic (red), rare population (yellow), intermediate population (orange), common population (green), and benign (blue). (*Right*) Pairwise KS statistics for RSA; higher values indicate greater distributional divergence. (b) (*Left*) Fraction of variants annotated with intramolecular van der Waals (VdW) interactions across the five datasets. (*Right*) Pairwise normalized Cohen’s w values quantifying distributional divergence in VdW interaction annotation between datasets. (c) Mean KS statistics averaged across all continuous protein features (*left*) and mean normalized Cohen’s w averaged across all binary and categorical protein features (*right*), summarizing distributional differences between variant datasets. See **Supplementary** Fig. 1 for all individual features. (d) Forest plots of odds ratios (OR, 95% CI) for representative protein features showing mild OR amplification as the control group shifts from rare to common population variants: α-helix, domain, and region/disordered. OR > 1 (red) indicates enrichment in pathogenic variants; OR < 1 (blue) indicates depletion. (e) Forest plots for representative features showing dramatic OR amplification or signal that emerges only when common population variants are used as the control: active site, intramolecular disulfide bond, and signal peptide. (f) Forest plot for all 103 protein features computed using the combined control group of benign and common population variants, organized across six feature attributes. The heatmap (*top*) shows OR values for all pairwise amino acid physicochemical group substitution along with the FDR-corrected p-values, i.e. q-values. Gray squares indicate features that did not reach statistical significance (*q* < 0.01).

To further evaluate how the choice of the control group from population variants against pathogenic variants affects the characterization of pathogenic variants, we computed the enrichment of pathogenic variants (odds ratio, OR) compared to each gnomAD allele frequency subgroup across all protein features (**Supplementary Fig. 2a**). For most of significantly enriched features, their effect sizes increase monotonically as the control group shifts from rare to common variants (**Supplementary Figs. 2b** and **2c**). **Figs. 2d** and **2e** highlight representative features illustrating two distinct patterns of OR amplification. For features showing mild but consistent OR amplification (**Fig. 2d**), some of the features, such as α-helices or domain annotation (from UniProtKB), show modest increases in OR (α-helix: 1.31 to 1.46; Domain: 1.99 to 2.38), while disordered region annotation (from UniProtKB) shows a modest decrease (0.19 to 0.17). The consistent direction and similar effect sizes of enrichment across rare, intermediate, and common population variants highlight that these features robustly differentiate pathogenic from population variants regardless of the choice of control group. In contrast, features with dramatically amplified OR when using common variants as the control (**Fig. 2e**), such as active sites (8.97 to 67.11) and intramolecular disulfide bonds (14.92 to 71.09) show that their discriminative power is highly dependent on the choice of control group. Some features, such as signal peptide annotation, showed near-neutral OR against rare variants but gained signal when common variants were used as control, indicating these features are only selectively depleted from common variants.

Collectively, these results support the use of common population variants as effective proxy-benign controls. For the rest of the paper, we used a combined control group of 25,871 common population variants and 104,848 benign variants, comprising 130,719 variants in total. **Fig. 2f** shows the OR of all 103 protein features from six protein feature groups using this combined control group.

For physicochemical properties, substitutions within the same physicochemical group (diagonal in the heatmap) are generally depleted in pathogenic variants, reflecting their tolerated nature. The exception is the *Special* group (Gly, Cys, Pro), whose members each perform structurally unique roles – cysteine forms disulfide bonds, glycine provides backbone flexibility, and proline restricts backbone – making even intra- group substitutions potentially disruptive. Consistently from the Grantham’s distance (*D*), severe amino acid substitutions (*D>150*) are enriched (OR=2.18) while mild changes (*D<50*) are depleted (OR=0.51) among pathogenic variants. Among structural features, pathogenic variants are enriched at core and buried positions (OR=3.45 and 1.82) and at intramolecular interaction sites, particularly disulfide bonds (OR = 14.99), while depleted at solvent-exposed (OR=0.29) and disordered regions (OR=0.19). Expectedly, structured residues of helices (3_10_, α, ρε; OR=1.23, 1.31, 2.33, respectively) and ý-bridge and sheets (OR=1.65 and 1.65) were found enriched among pathogenic variants and relatively unstructured polyproline helix and loops were depleted (OR=0.90 and 0.49).

Functional sites showed strong enrichment across all individual features (e.g. Active sites; OR = 9.01, binding sites; OR = 7.16). From the domain/region group, intramembrane regions were notably enriched (OR = 14.95) while features that have a connection to the disordered regions of proteins were depleted (Region/Disordered; OR = 0.19, Compositional bias; OR = 0.44). Among post-translational modifications, lipidation, acetylation, and methylation sites were moderately enriched (OR=2.32, 1.62, and 1.67, respectively), consistent with their roles at functionally critical positions. In contrast, phosphorylation, ubiquitination, and O-GalNAc/GlcNAc sites were depleted. Phosphorylation sites (OR=0.79) frequently occur in disordered or flexible regions (*47, 48*) and are often functionally redundant within signaling pathways, making single-site variants less likely to be strictly pathogenic. Ubiquitination (OR=0.84) targets proteins for degradation rather than marking positions of core functional importance, and glycosylation sites (O-GalNAc/GlcNAc; OR=0.41) tend to occur on solvent-exposed surfaces (*49*) where substitutions are generally better tolerated. Finally, all PPI features were strongly enriched, with intermolecular disulfide bonds showing the highest OR across the entire feature set (OR = 19.25), reflecting the critical importance of interface residues for protein function. These enrichment patterns directly inform PFES calculation, where each feature’s association with pathogenic variation contributes to the final score.

### Protein feature enrichment patterns of variants vary across functional classes

The enrichment analyses described above were performed across all human proteins collectively to identify protein features broadly impacted by pathogenic missense variants. However, proteins are functionally diverse, and the protein features impacting pathogenicity are unlikely to be uniform across all functional contexts (*18*). To address this, we repeated the enrichment analysis separately for 20 different protein functional classes (**Fig. 3a** and **Supplementary Table 1**) for the pathogenic and final control (benign and common population) datasets. PANTHER database (*38*) annotates proteins into 24 functional classes: 64% of all human proteins are annotated, while the remaining 34% are unclassified (**Methods**: Protein class annotation from PANTHER). **Supplementary Fig. 3** shows the number of case and control variants for the statistical analysis within each protein class. For the robustness of the statistical test, we merged 5 protein classes having fewer than 300 variants in either pathogenic or control datasets into an *unclassified* class, resulting in a total of 20 protein classes.

**Fig. 3.**
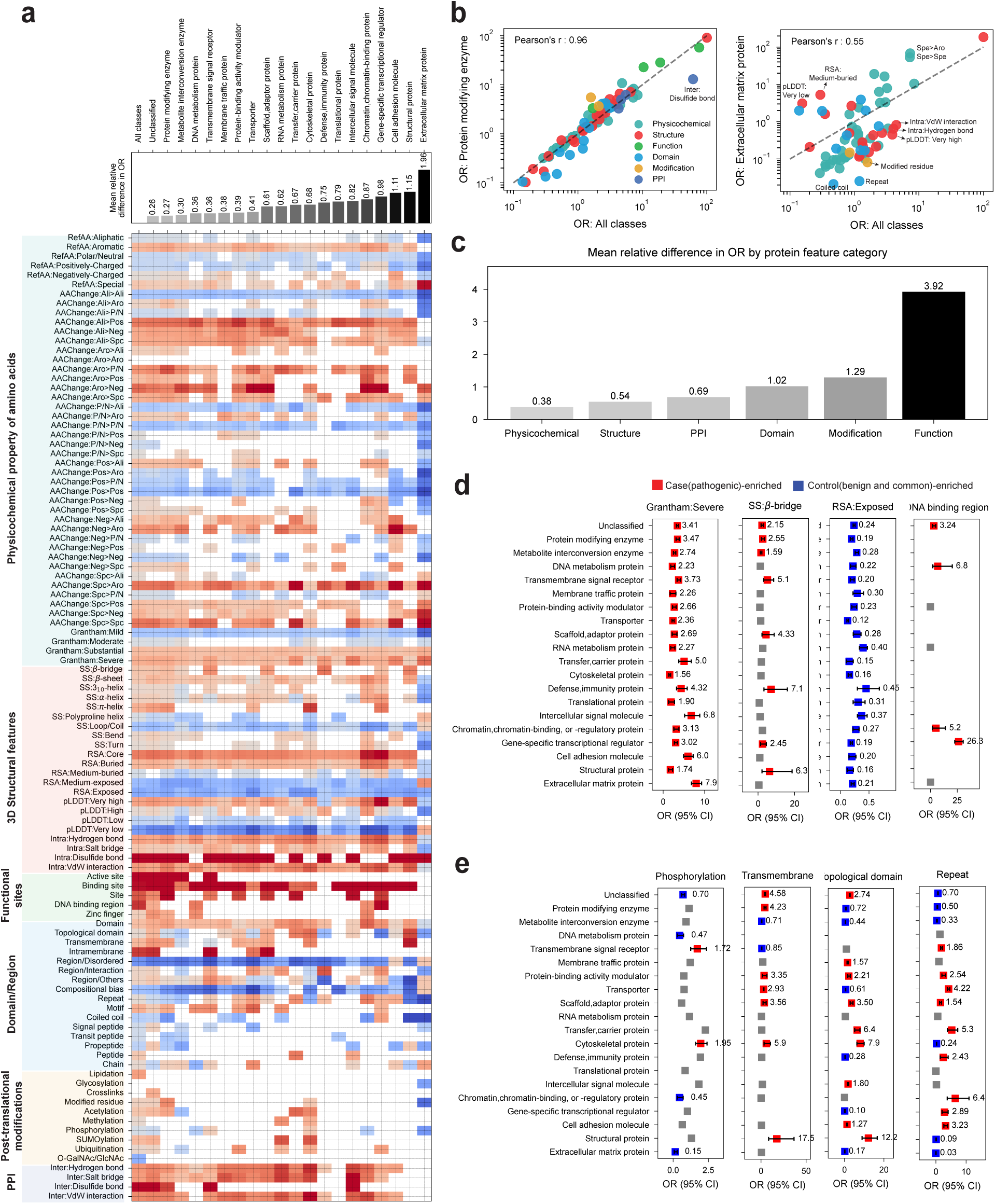
Protein feature enrichment patterns differ across protein’s functional classes. (a) (*top*) The mean relative differences in class-specific ORs from the all-classes-averaged baseline. (*bottom*) Heatmap showing the enrichment of all protein features across 20 PANTHER-defined protein classes. Each cell represents the odds ratios (OR) from Fisher’s exact test comparing pathogenic versus control variants within each protein class. Red indicates features enriched in pathogenic variants (OR > 1), blue indicates features enriched in control variants (OR < 1), with color intensity reflecting the magnitude of enrichment. White cells indicate either no enrichment or features that did not reach statistical significance (FDR-corrected *p* < 0.01). **(b)** Per-feature OR values calculated within a protein class are plotted against the corresponding all-classes OR. The least divergent classes (protein-modifying enzyme) and the most divergent classes (extracellular matrix protein) are shown. Each point represents an individual feature, colored by feature group. The dashed diagonal line indicates perfect agreement with the all-classes estimate; points above or below the diagonal indicate amplified or weakened enrichment relative to the proteome-wide estimate, respectively. **(c)** Mean relative difference in class-specific ORs from the all-classes-averaged baseline, summarized by protein feature categories. Each bar represents the average deviation across all protein classes within each feature category, illustrating that functional site annotations show the greatest class-dependent variability while physicochemical properties show the least. Forest plots of odds ratios (OR, 95% CI) for representative features are given: **(d)** Severe Grantham’s distance, SS:β-bridge, RSA:Exposed, and DNA binding region, where they showed consistent enrichment direction but variable effect sizes across classes, and **(e)** Phosphorylation, Transmembrane, Topological domain, and Repeat, where they showed enrichment direction inverted depending on protein class context. Grey squares indicate features that did not reach statistical significance (FDR-corrected *p* < 0.01) within a given protein class.

Protein class-specific analysis revealed that while the broad direction of feature enrichment (OR) is largely preserved across classes, the magnitude varied substantially depending on protein class identity (**Fig. 3**). To quantify this, we calculated the mean relative difference in OR for each class from the all-classes baseline, i.e., (OR_class_ – OR_All classes_))/OR_All classes_ (*upper*, **Fig. 3a**). Values ranged from near-baseline for unclassified proteins (0.26, i.e. 26% difference) and protein-modifying enzyme (0.27), indicating that their pathogenic variant enrichment patterns are similar to all proteins combined), to markdly evevated for structural protein (1.15) and extracellular matrix protein (1.96), indicating highly divergent, class-specific feature enrichment patterns. We further plotted per-feature OR values calculated within protein-modifying enzymes and extracellular matrix proteins, the least and the most divergent classes, respectively, against all-classes OR values (**Fig. 3b**). Protein-modifying enzymes showed tight clustering along the diagonal across nearly all feature categories, while extracellular matrix proteins showed a high degree of divergence from the proteome-wide protein feature profile. This divergence was most pronounced across 3D structural features, including confidence-extreme pLDDT values (very low and very high), medium-buried RSA, and intramolecular VdW interactions and hydrogen bonds, but extended to other protein feature categories as well, such as modified residue and repeat annotations.

To identify which feature attributes drive this class-dependent variation, we decomposed the mean relative difference by protein feature groups. **Fig. 3c** shows that functional site annotations exhibited the greatest class-dependent variability, followed by modification-associated and domain/region annotations, while physicochemical properties exhibited the smallest deviation across classes. Physicochemical properties of amino acid substitutions are relatively universal and broadly applicable regardless of protein function. Functional site annotation (active site, DNA binding region, etc.), by contrast, is inherently tied to the specific roles of proteins in a family (e.g., gene-specific transcriptional regulators or extracellular matrix proteins), and therefore represents a class-specific feature enriched among pathogenic variants.

Examining individual features across all protein classes revealed two qualitatively distinct patterns of class-dependent behavior. The first encompasses features that are enriched in the same direction across nearly all classes but with substantially different effect sizes (**Fig. 3d**). Severe Grantham’s distance and β-bridge were consistently enriched among pathogenic variants across all classes, though effect sizes varied considerably, e.g., β-bridge enrichment ranged from OR = 1.59 in metabolite interconversion enzymes to OR = 7.1 in defense/immunity proteins. Surface-exposed residues were consistently depleted across all classes. DNA-binding region enrichment was the most extreme example of class-restricted directionality, reaching OR = 26.3 in gene-specific transcriptional regulators and OR = 6.8 in DNA metabolism proteins, with no significant enrichment in any other class. The second pattern comprises features whose enrichment direction inverts across classes (**Fig. 3e**). Phosphorylation sites were enriched in transmembrane signal receptors (OR = 1.72) and cytoskeletal proteins (OR = 1.95) but depleted in extracellular matrix proteins (OR = 0.15) and chromatin-binding proteins (OR = 0.45), with no significant signal in the remaining classes. Transmembrane, topological domain, and repeat annotations showed similarly class-restricted patterns, with strong enrichment in classes where these properties define protein architecture, i.e. structural proteins (transmembrane OR = 17.5; topological domain OR = 12.2), and depletion or absence elsewhere. In each case, significant enrichment was observed only in classes where the annotated feature is a defining structural or functional property.

### PFES provides molecular context-based interpretation of pathogenic variants

Having established that pathogenic variants exhibit distinct, protein class-specific enrichment patterns across protein features defining their molecular context, we developed a Protein Feature Enrichment Score (PFES; **Fig. 1d** and **Methods**: Computation of Protein Feature Enrichment Score (PFES)). PFES quantifies the degree to which a missense variant displays a protein functional class-specific molecular context statistically associated with pathogenic or benign variations. It is calculated as the sum of log-odds ratios across significantly enriched protein features, and decomposed into contributions from each protein feature group: physicochemical properties of amino acids (PFES_Physicochemical_), 3D structural features (PFES_Structure_), domain/region (PFES_Domain_), functional sites (PFES_Function_), post-translational modifications (PFES_Modification_), and protein-protein interactions (PFES_PPI_).

**Fig. 4a** shows the empirical PFES distributions across pathogenic, control (benign and common population), and VUS datasets. Pathogenic variants shift toward positive values, and control variants concentrate near zero or negative values, but the two distributions overlap substantially. This overlap is biologically expected: variants acting through regulatory perturbations, altered expression, or haploinsufficiency may lack a broad protein feature enrichment signature, and the overlap reflects the mechanistic heterogeneity of pathogenic variation rather than classification noise. VUS occupy an intermediate position consistent with their unresolved clinical status, and their distribution across protein feature (PF) categories provides a molecular basis for prioritization that we examine in detail in the later section.

**Fig. 4.**
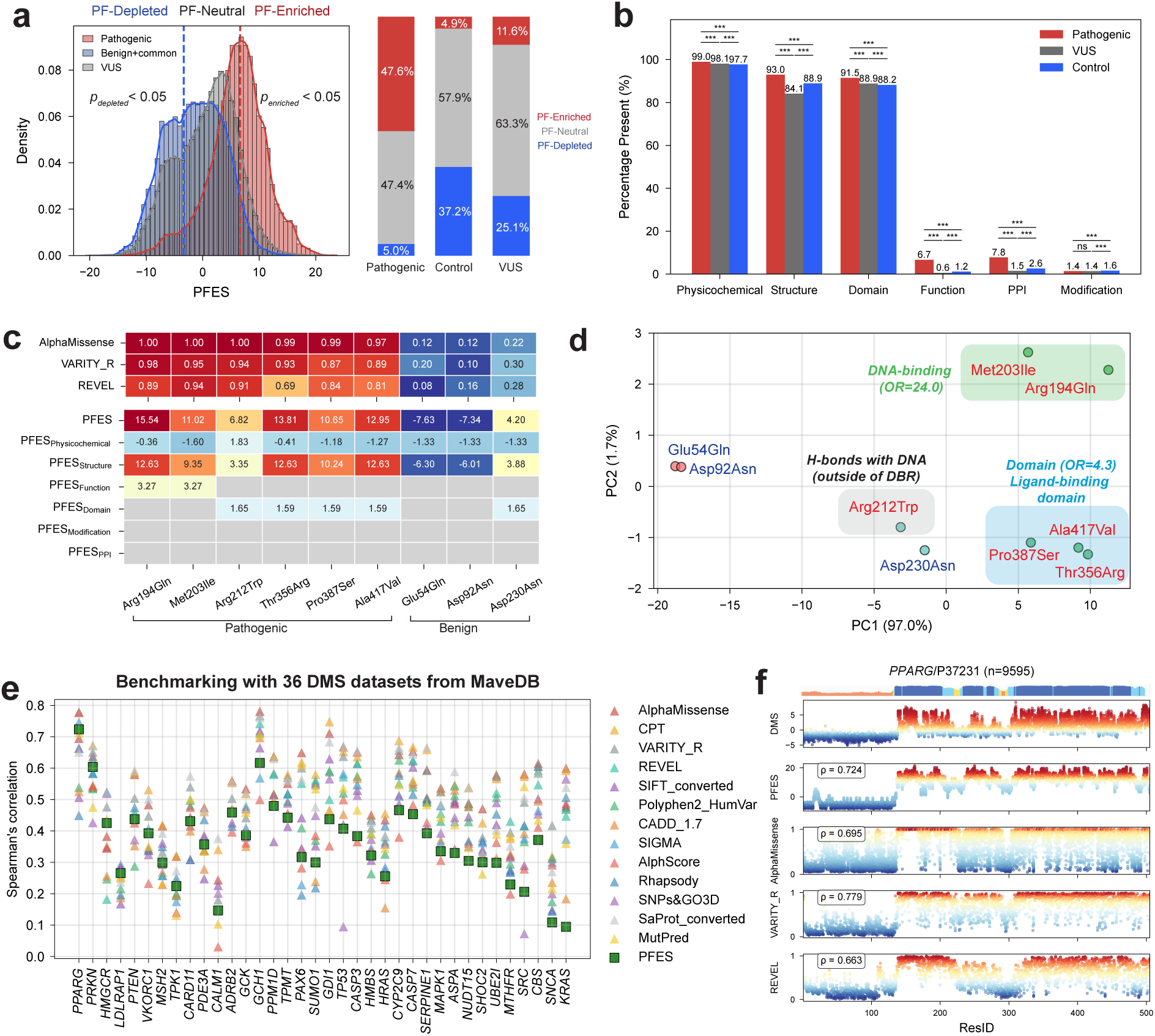
Mechanistic partitioning of missense variants by PFES and benchmarking against deep mutational scanning data. **(a)** (*Left*) Empirical PFES distributions for pathogenic, control, and VUS datasets. Dashed lines indicate PF-Enriched and PF-Depleted thresholds derived from statistical deviation from each distributions (*p_enriched_* or *p_depleted_* < 0.05; one-sided p-values. See **Methods:** for more details). (*Right*) Stacked bar charts showing the proportion of PF-Enriched, PF-Neutral, and PF-Depleted variants within each dataset. **(b)** Percentage of variants presenting each score attribute from protein feature groups across pathogenic, VUS, and control (benign+common) datasets. Significance brackets indicate pairwise comparisons between pathogenic and control variants by pairwise chi-square *p*-values (* *p* < 0.05, ** *p* < 0.01, *** *p* < 0.001, ns: not significant). **(c)** Heatmap of VEP scores and PFES and its subscores for 6 pathogenic and 3 benign *PPARG* variants from Majithia *et al.* (*50*). **(d)** Principal component analysis (PCA) of 9 *PPARG* variants based on six subscores. Shaded areas indicate variant clusters and contributing protein features with odds ratios (OR) are given. **(e)** Spearman’s correlation between PFES and 13 VEPs against 36 DMS datasets. Each point represents one DMS dataset; PFES is shown as green squares (**see Supplementary Table 2** for full data) **(f)** Residue-level score profiles for *PPARG*/P37231 comparing DMS, PFES, AlphaMissense, VARITY_R, and REVEL along the protein sequence. Spearman’s correlation values against DMS scores are shown for each VEP and PFES. Points are colored by each score magnitude.

We partitioned variants into three statistically defined categories based on their position relative to the empirical pathogenic and control distributions (**Methods:** PFES-based partitioning of PF-Enriched, Neutral, and Depleted variants). Among pathogenic variants, 48% were PF-Enriched, enrichment for protein features associated with pathogenicity, 47% PF-Neutral, and 5% PF-Depleted (**Fig. 4a**). Among controls, only 37% were PF-Depleted, showing enrichment for benign-associated features such as surface-exposed or disordered regions, while more than half (58%) of variants remained PF-Neutral. This asymmetry in the population of partitioned variants between pathogenic and control datasets indicates that pathogenic variants tend to produce distinct, recognizable molecular signatures, whereas tolerance is often characterized by the absence of distinctive protein feature signals rather than the presence of specific tolerance-promoting characteristics. The VUS distribution provides critical insight into the utility of feature enrichment analysis for variants of uncertain significance. Aligned with most VUS often lacking strong disruptive protein feature signals, the majority of VUS were PF-Neutral (63%) and smaller fractions of PF-Depleted (25%) and PF-Enriched (12%) VUS. The 12% of PF-Enriched VUS represent variants warranting prioritization for functional studies or additional clinical investigation, as they exhibit protein feature profiles statistically consistent with known pathogenic variants. Conversely, the 25% of PF-Depleted VUS may represent variants more likely to be reclassified as benign upon additional evidence.

Examining contributions from different feature attributes provided further insight into how individual feature groups shape the overall PFES signal, as measured by the percentage of variants presenting a score contribution from each attribute category (**Fig. 4b**). A variant may lack a subscore for a given feature attribute if it has no annotated features within that group or none of the features reach statistical significance. Subcores from physicochemical properties, structural, and domain feature groups were present in the majority of variants (>84%) across all three datasets, and their subscore distributions were well-separated between pathogenic and control variants (**Supplementary Fig. 4**), indicating that their magnitude and sign carry discriminating information. Functional and PPI features were selectively annotated but strongly enriched in pathogenic variants relative to controls (Function: 6.7% vs. 1.2%; PPI: 7.8% vs. 2.6%), and their scores were consistently positive, suggesting that their primary signal lies in presence rather than magnitude. Modification features showed a distinct pattern: annotation rates were low and comparable across variant datasets (less than 2%), but score distributions showed moderate bidirectional separation between pathogenic and control variants, indicating that when present, the direction of modification-associated change carries interpretive value. Together, these results highlight that protein features contribute to pathogenicity through multiple modes, reflecting not only the degree of physicochemical and structural disruption, but also the involvement of functionally critical, protein-specific features whose perturbation is associated with pathogenicity.

To demonstrate the interpretive value of PFES decomposition in a real clinical genetics context, we examined a set of *PPARG* variants reported by Majithia *et al.* (*50*), in which all possible *PPARG* missense variants were evaluated using a pooled functional assay to train a pathogenicity predictor, resulting in 6 novel pathogenic (Arg194Gln, Met203Ile, Arg212Trp, Thr356Arg, Pro387Ser, and Ala417Val) and 3 benign (Glu54Gln, Asp92Asn, and Asp230Asn) variants. **Fig. 4c** shows the VEP scores (AlphaMissense, VARITY_R, and REVEL), where they discriminate pathogenic and benign variants, and the PFES and subscores for those variants. PFES showed that all 6 pathogenic variants fell within the PF-Enriched range, while 2 out of 3 benign variants (Glu54Gln and Asp92As) partitioned into the PF-Depleted range (Asp230Asn; PF-Neutral). Decomposed subscores further showed that structural features dominated the pathogenic signal across most variants, while function and domain features were selectively presented. The principal component analysis (PCA) on the 6 subscores separates variants into mechanistically interpretable clusters (**Fig. 4d**). Arg194Gln and Met203Ile clustered together, attributed by DNA-binding site annotation (OR=24.0); Thr356Arg, Pro387Ser, and Ala417Val formed a separate cluster with the ligand-binding domain annotation (OR=4.3). Benign variants (Glu54Gln and Asp92Asn) showed negative PFES_Structure_ and PFES_Physicochemical_, clustering outside both pathogenic groups. Arg212Trp occupied an intermediate position between the two pathogenic clusters, spatially distinct from both, hinting at a mechanism that does not align cleanly with either canonical pathogenic context.

Arg212Trp illustrates the hypothesis-generating potential of this decomposition approach. While its overall PFES correctly classifies it as pathogenic, its decomposed subscore profiles show atypically low PFES_Structure_ compared to all other pathogenic variants, with only a modest physicochemical contribution. Existing VEPs provide no indication that Arg212Trp is mechanistically distinct: AlphaMissense (1.00), VARITY_R (0.94) and REVEL (0.91) score it as confidently pathogenic and essentially indistinguishable from other pathogenic variants, for example, Met203Ile (AlphaMissense; 1.00, VARITY_R; 0.95, and REVEL; 0.94). The atypical subscore profile prompted further structural investigation, which revealed that Arg212 directly contacts DNA through hydrogen bonding outside the canonically annotated DNA-binding region, a mechanism not captured within current PFES annotations. This case illustrates that atypical attribute profiles can flag variants whose pathogenic mechanism requires deeper investigation, generating testable hypotheses even at the boundaries of current annotation coverage.

### Benchmarking of PFES with functional readouts from MaveDB

The landscape of variant effect prediction has evolved dramatically with a massive number of available variants and advanced deep learning approaches, with many state-of-the-art predictors achieving remarkable accuracy. To contextualize PFES within the broader landscape of variant effect predictors (VEP), we benchmarked its performance against 13 VEPs using 36 deep mutational scanning (DMS) datasets (*32, 51*) using Spearman’s correlation between predicted scores and experimental readouts (**Fig. 4e** and **Supplementary Table 2; Methods**: Benchmarking with deep mutational scanning data and variant effect predictors). The 13 VEPs were selected to represent the diversity of prediction approaches: deep learning methods (AlphaMissense (*30*), CPT-1 (*52*), VARITY_R (*25*)), a metapredictor (REVEL (*29*)), traditional tools (SIFT (*20*), PolyPhen-2 (*24*), CADD (*28*)), and predictors that leverage protein’s 3D structures (SIGMA (*53*), AlphScore (*45*), Rhapsody (*54*), SNPs&GO3D (*27*), SaProt (*26*), MutPred (*55*)). It is important to note that PFES was not optimized or calibrated as a VEP to achieve high predictive accuracy. Instead, this benchmarking shows how the molecular context of mutations quantified through PFES provides mechanistic interpretation of variants and DMS data.

Across 36 DMS datasets, no tool emerged as a universal winner: performance varied substantially across genes for all predictors, including PFES. Despite no direct optimization toward DMS functional readouts, PFES showed reasonable agreement across many datasets, performing well for some genes and more modestly for others. As an illustrative example, *PPARG* showed strong spatial concordance between DMS and PFES score profiles along the sequence (**Figure 4f**), with PFES achieving ρ = 0.724, comparable to AlphaMissense (ρ = 0.695) and REVEL (ρ = 0.663). The agreement reflects not only numerical correlation but spatial coherence – both DMS and PFES identify the same constrained regions along the protein sequence, which would be critical to its function.

The decomposed subscore profiles provide mechanistic insight into why PFES captures the *PPARG* mutational landscape effectively (**Supplementary Figure 5**). Structural features drive the broad score elevation across the functional regions of the protein (residues 130-210 and 300-500), while functional annotations (DNA-binding and ligand-binding sites) and domain features (region/interaction and lipid-binding domain annotations) explain the specific hotspot regions where both DMS and PFES scores are highest. PPI and modification features contribute selectively at residues carrying the relevant annotations, further refining the score landscape. This attribute-resolved view reveals what other VEPs cannot provide: not only where the hotspot is, but which specific protein characteristics underlie its functional sensitivity.

The benchmarking performance of PFES is further influenced by a subtler factor: the sequence coverage of individual DMS datasets. Some proteins in MaveDB are represented by multiple DMS experiments covering different sequence ranges, and sometimes it covers only a functionally important sub-region. Because PFES is designed to capture the global mutational tolerance landscape across the full protein sequence, identifying constrained versus unconstrained regions, its correlation with DMS is inherently reduced when the assay already focuses exclusively on a high-constraint region, leaving little landscape variation for the score to track. *KRAS* and *TP53* illustrate this effect clearly (**Supplementary Figure 6**). When evaluated against domain-focused DMS datasets, PFES correlations were modest (KRAS: ρ = 0.094; TP53: ρ = 0.407). Switching to full-length DMS datasets covering the entire protein sequence substantially improved performance for both proteins (*KRAS*: ρ = 0.198; *TP53*: ρ = 0.611), with *TP53* PFES correlation surpassing AlphaMissense (ρ = 0.512), REVEL (ρ = 0.524), and CADD (ρ = 0.436). These results suggest that PFES is particularly well-suited for capturing global mutational sensitivity landscapes, and that benchmarking against partial-coverage DMS datasets may underestimate its performance. Collectively, these results suggest that DMS coverage is an important consideration when benchmarking any proteome-wide scoring method, and that full-length experimental assays provide a more appropriate evaluation framework for tools designed to capture global mutational landscapes.

### Prioritization and triage of variants of uncertain significance by the PFES framework

A central challenge in clinical genetics is the growing burden of variants of uncertain significance (VUS), which lack sufficient evidence for pathogenic or benign classification. We reasoned that PF-Enriched VUS – those exhibiting molecular context defined by pathogenic variant-associated protein features– represent high-priority candidates for further functional investigation. To evaluate this utility, we applied PFES-based partitioning to a proteome-wide VUS dataset curated from ClinVar.

Across the human proteome, PF-Enriched VUS were widely distributed, with most genes (n=9,154) harboring fewer than ten such variants, while a smaller subset exhibited substantially higher counts (over 201 VUSes; n=75) (**Fig. 5a**). Notably, the ten genes with the greatest number of PF-Enriched VUS, including *LZTR1*, *ATM,* and *CFTR*, are well-established rare disease genes (**Fig. 5b**). This convergence confirms that PFES selectively flags gene variants under strong protein structure–function constraint and identifies genes in which VUS triage carries the greatest clinical consequence.

**Fig. 5.**
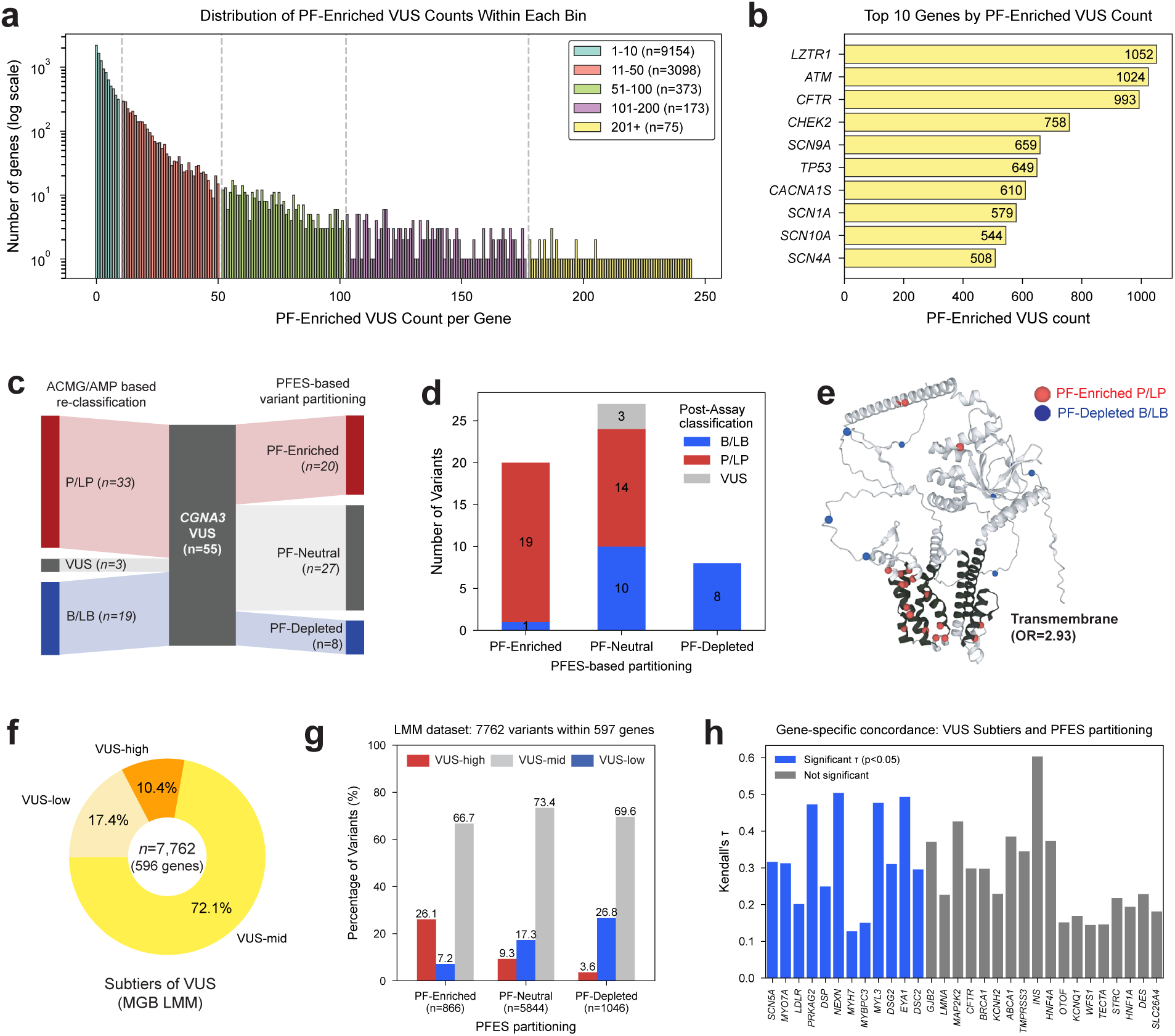
PFES-based partitioning enables prioritization and triage of variants of uncertain significance. **(a)** Distribution of PF-Enriched VUS counts per gene across the human proteome (log scale), colored by count bin. **(b)** Top 10 genes ranked by PF-Enriched VUS count. Genes are highlighted in yellow, reflecting their designation as established rare disease genes in the GenCC database. **(c)** Re-classification of 55 *CNGA3* missense VUS from functional assay by Solaki *et al.* (*12*) (*left*) alongside their PFES-based partitioning (*right*). Numbers indicate variant counts per category. **(d)** Stacked bar chart comparing post-assay reclassification of variants (B/LB, P/LP, VUS) across PFES-based partitioning. Numbers within bars indicate variant counts. **(e)** Mapping of concordant variants onto the *CNGA3* AlphaFold2 structure. PF-Enriched P/LP variants (red) and PF-Depleted B/LB variants (blue) are shown as spheres. **(f)** Donut chart showing the population of the VUS subtiers (VUS-low, VUS-mid, VUS-high) made from Mass General Brigham Laboratory of Molecular Medicine (MGB LMM). **(g)** Percentage of VUS subtiers across PF-Enriched, PF-Neutral, and PF-Depleted variants. **(h)** Kendall’s τ to measure concordance between VUS subtier assignment and PFES partitioning across 596 genes, ranked by −log(*p*-value). Blue bars indicate genes with statistically significant positive concordance (*p*<0.05; *n*=14); gray bars indicate non-significant associations.

To assess whether PF-Enriched variants reflect genuine pathogenic potential, we examined a published functional screening dataset for *CNGA3* by Solaki *et al.* (*12*), a cyclic nucleotide-gated channel subunit associated with achromatopsia, in which missense VUS were reclassified according to ACMG/AMP guidelines based on experimental readouts. Of 55 missense VUS in *CNGA3*, the addition of functional readout to existing evidence reclassified 19 as B/LB, 33 as P/LP, and 3 remained as VUS (**Fig. 5c**). PFES partitioning independently identified 20 as PF-Enriched, 27 as PF-Neutral, and 8 as PF-Depleted. Comparing these categories against functional reclassification revealed a striking concordance: PF-Enriched variants were predominantly reclassified as P/LP (19 of 20, 95%), and all of PF-Depleted variants were reclassified as B/LB (8 of 8, 100%) (**Fig. 5d**). Structural visualization further confirmed that PF-Enriched P/LP variants cluster at functionally critical positions, including the transmembrane domain (OR=2.93), whereas PF-Depleted B/LB variants map to structurally disordered regions (**Fig. 5e**). These results demonstrate that PFES-based partitioning captures functionally meaningful variation in *CNGA3*, where protein feature enrichment aligns with experimentally validated pathogenic consequence, establishing a basis for large-scale evaluation across the broader clinical VUS landscape.

To extend this validation to a larger and more diverse clinical context, we applied PFES-based partitioning to a VUS dataset from Mass General Brigham Laboratory for Molecular Medicine (MGB LMM) (**Fig. 5f**) in which VUSes were subclassified into three subtiers based on the similarity to benign variation, i.e., VUS-high, VUS-mid, and VUS-low(*56*) (7,762 variants among 596 genes). The majority of variants fell into VUS-mid (72.1%), with VUS-high and VUS-low comprising 10.4% and 17.4%, respectively. VUS partitioned into PF-Enriched showed the highest fraction of VUS-high (26.1%), while only 3.6% of PF-Depleted VUS were VUS-high. Similarly, the fraction of VUS-low progressively decreased from PF-Depleted VUS (26.8%) to PF-Enriched VUS (7.2%). This is consistent with the expectation that pathogenicity-leaning VUS, i.e., VUS-high, are more likely to exhibit protein features associated with pathogenicity, thus likely to be partitioned into PF-Enriched. To quantify gene-level concordance between VUS subtier and PFES partitioning, we calculated Kendall’s rank correlation coefficient (τ) for each gene represented in the dataset (**Methods**: Evaluation of statistical concordance). Rather than uniform concordance across all genes, we observed meaningful heterogeneity: 14 out of 596 genes showed statistically significant positive concordance (*p*<0.05), while others exhibited weaker or non-significant associations (**Fig. 5h**). This gene-level variability is itself informative: genes with strong concordance likely harbor variants whose pathogenicity is mediated primarily through disruption of protein features directly captured by PFES, whereas genes with weaker concordance may involve more context-specific or gain-of-function mechanisms that operate outside this framework.

To illustrate this concretely, we examined two genes with significant Kendall’s τ: *SCN5A* and *PRKAG2* (**Supplementary Fig. 7**). In *SCN5A*, PF-Enriched variants had the highest fraction of VUS-high (22%), mapped predominantly to the transmembrane domain, while PF-Depleted variants possessed no VUS-high but 36% of VUS-low. *PRKAG2* exhibited an even more pronounced pattern despite a smaller sample size: the single PF-Enriched variant was classified as VUS-high, while 45% of PF-Depleted variants were VUS-low. Together, these gene-level examples illustrate how PFES partitioning recapitulates clinical subclassification judgment from first principles of protein feature enrichment, and how structural context reinforces the mechanistic interpretation of each partition category. Collectively, these analyses demonstrate that PFES-based partitioning provides actionable molecular stratification of VUS, with utility both for prioritizing variants for functional follow-up and for interpreting subclassification decisions in a clinical setting.

### Decomposed PFES attributes differentiate loss- and gain-of-function variants

Pathogenic missense variants can drive disease through distinct molecular mechanisms: Loss-of-function (LoF) variants, typically reducing or abolishing normal protein activity through destabilization, misfolding, or disruption of catalytic and binding functions (*57–59*), and gain-of-function (GoF) variants, showing aberrant or enhanced activity, often through constitutive activation, dominant negative effects, or acquisition of novel protein-protein interactions (*60*). Distinguishing LoF and GoF variants carries clinical relevance and inform treatment strategies, inheritance patterns, and the interpretation of de novo variant, however, existing VEPs are trained to separate pathogenic from benign variation without mechanistic distinction. We hypothesize that the different molecular context of LoF/GoF variants can be captured by PFES and protein feature-based interpretation. To test this, we assembled a dataset of 1,284 LoF, 1,753 GoF, and 4,097 control variants (**Fig. 6a**) from Gene2Phenotype (*61*) across 176 genes (**Supplementary Table 3** and **Methods**: Collection of Loss-and Gain-of-function variants from Gene2Phenotype) and addressed three questions: whether PFES distinguishes pathogenic from benign variation regardless of mechanism; whether protein feature composition differs between LoF and GoF variants; and whether any mechanism-associated feature patterns are consistent across genes.

**Fig. 6.**
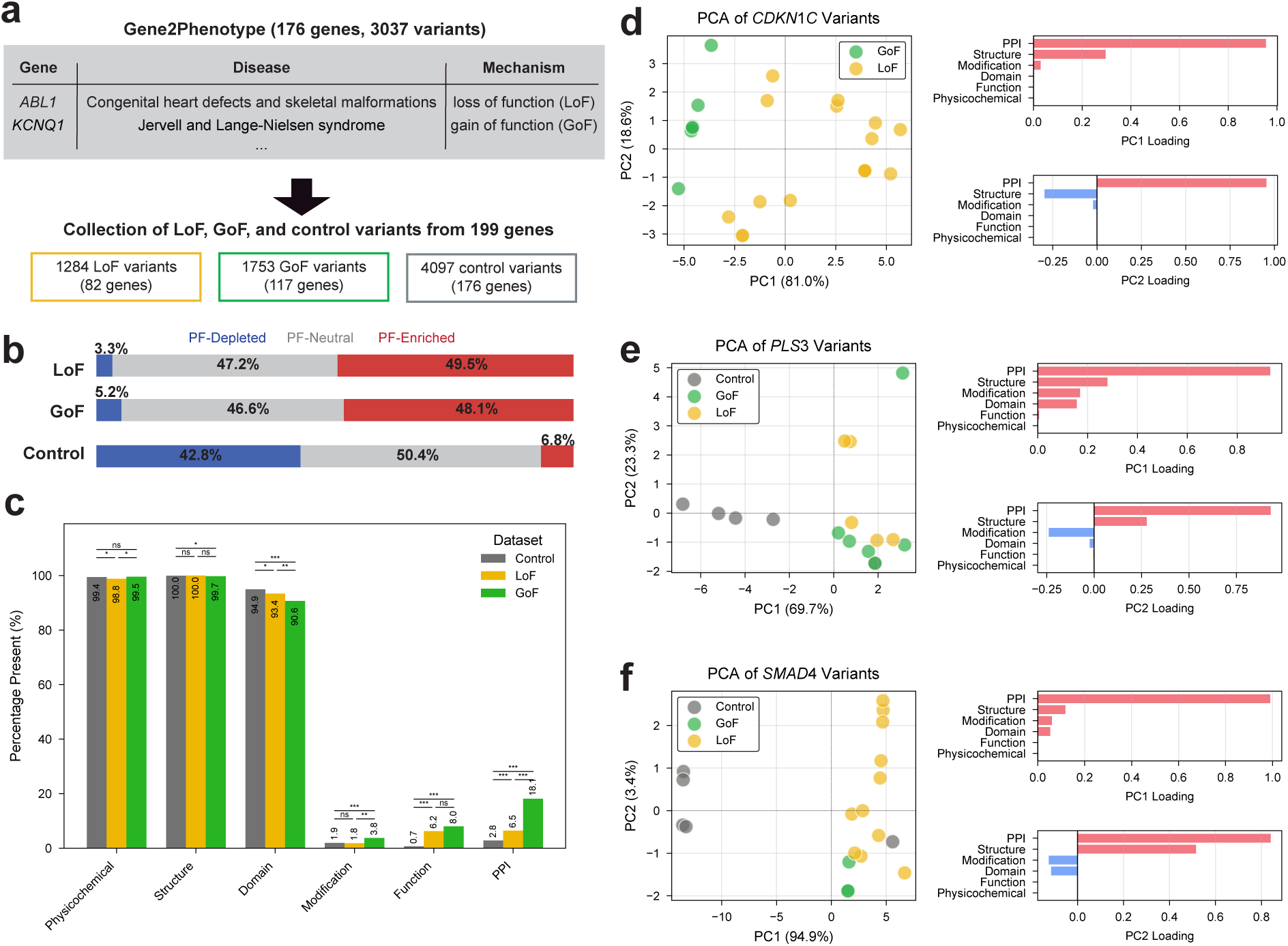
PFES captures pathogenic protein feature disruption across loss- and gain-of-function mechanisms. **(a)** Variants were curated from Gene2Phenotype based on annotated disease mechanism, yielding 1,284 loss-of-function (LoF), 1,753 gain-of-function (GoF), and 4,097 control variants across a broad range of Mendelian disease genes (See **Supplementary Table 3** for the full table of gene, disease, mechanism annotation). **(b)** Distribution of PFES-based partitioning categories (PF-Enriched, PF-Neutral, PF-Depleted) across LoF, GoF, and control variant groups. Both LoF and GoF variants are predominantly PF-Enriched, while control variants are predominantly PF-Depleted. **(c)** Percentage of variants presenting each subscore from six attributes across LoF, GoF, and control variants. Statistical significance between groups is indicated by *p*-values from a pairwise chi-square test (* *p* < 0.05, ** *p* < 0.01, *** *p* < 0.001, ns: not significant). **(d-f)** Principal component analysis (PCA) on six PFES subscores for variants from three exemplar genes: **(d)** *CDKN1C,* **(e)** *PLS3,* and **(f)** *SMAD4*. (*Left*) LoF, GoF, and control variants in the first and second PCA axes, with the percentage of variance explained indicated on each axis. (*Right*) The feature attribute loadings for PC1 (*top*) and PC2 (*bottom*). PPI features consistently show the highest PC1 loadings across all three genes, with GoF variants clustering toward positive PC1 values and LoF variants toward negative values.

PFES-based partitioning confirmed that both LoF and GoF variants were predominantly PF-Enriched (49.5% and 48.1%), while controls were predominantly PF-Depleted (42.8%), with few variants PF-Enriched (6.8%; **Fig 6b**). Existing VEPs similarly separate controls from LoF/GoF variants by score (**Supplementary Fig. 8**). In contrast, PFES can be decomposed into its protein feature attributes illuminating the molecular context driving the LOG/GoF mechanisms (**Fig. 6c**). While physicochemical, structural, and domain features were near-universally present across all three groups (around 90%), these categories alone offer limited discriminating power among LoF, GoF, and control variants. In contrast, modification (PTMs), function (active/binding sites), and PPI features were present at substantially lower frequencies but showed significant and informative differences between groups. PTMs were modestly but significantly enriched in GoF variants (3.8%) relative to both LoF (1.8%) and controls (1.9%). The most striking difference was observed for PPI features, which were nearly three times more prevalent in GoF variants (18.1%) than in LoF variants (6.5%), and both were significantly elevated above controls (2.8%). This disproportionate enrichment of PPI features in GoF variants suggests that they more frequently occur at protein interaction interfaces, consistent with pathogenic mechanisms involving aberrant complex formation, dominant negative effects, or neo-morphic interactions.

To examine how these feature attributes manifest at the gene level, we performed principal component analysis (PCA) on 6 subscores for three genes in which both LoF and GoF variants were present and led to distinct phenotypes (**Figs. 6d-f**). *CDKN1C* GoF variants cause IMAGe syndrome through increased protein stability driving growth restriction, while LoF variants cause Beckwith-Wiedemann syndrome through loss of cell cycle inhibition. *PLS3* GoF variants cause congenital diaphragmatic hernia through altered actin dynamics, while LoF variants cause X-linked osteoporosis through disrupted actin bundling. *SMAD4* GoF variants cause Myhre syndrome through dominant-negative effects, while LoF variants cause juvenile polyposis and hereditary hemorrhagic telangiectasia through haploinsufficiency. In all three genes, PC1 captured the substantial majority of variance (81.0%, 69.7%, and 94.9% for *CDKN1C*, *PLS3*, and *SMAD4*, respectively; **Figs. 6d-f**, *left*), and PPI features consistently showed the highest PC1 loadings (**Figs. 6d-f**, *right*). PC2, which captured secondary axes of variation (18.6%, 23.3%, and 3.4%, respectively), received contributions from structure and modification features, with some genes showing negative loadings for specific categories, indicating that these features contribute to opposing directions depending on the protein context. This gene-level heterogeneity underscores an important interpretive principle: proteome-wide trends in feature enrichment identify general associations between protein characteristics and disease mechanism, but the manifestation of each feature attribute is inherently gene-specific.

### PFES as a proteome-wide missense variant interpretation and gene prioritization resource

To support broad application of the PFES framework, we compiled PFES calculations and PFES-based partitioning across all 223 million possible missense substitutions in 20,232 human proteins corresponding to 20,153 genes (**Fig. 7a**). Proteome-wide, 17.6% of all possible substitutions are PF-Enriched, 61.9% PF-Neutral, and 20.5% PF-Depleted, establishing that the majority of theoretically possible amino acid changes lack a distinctive protein feature signal in either direction.

**Fig. 7.**
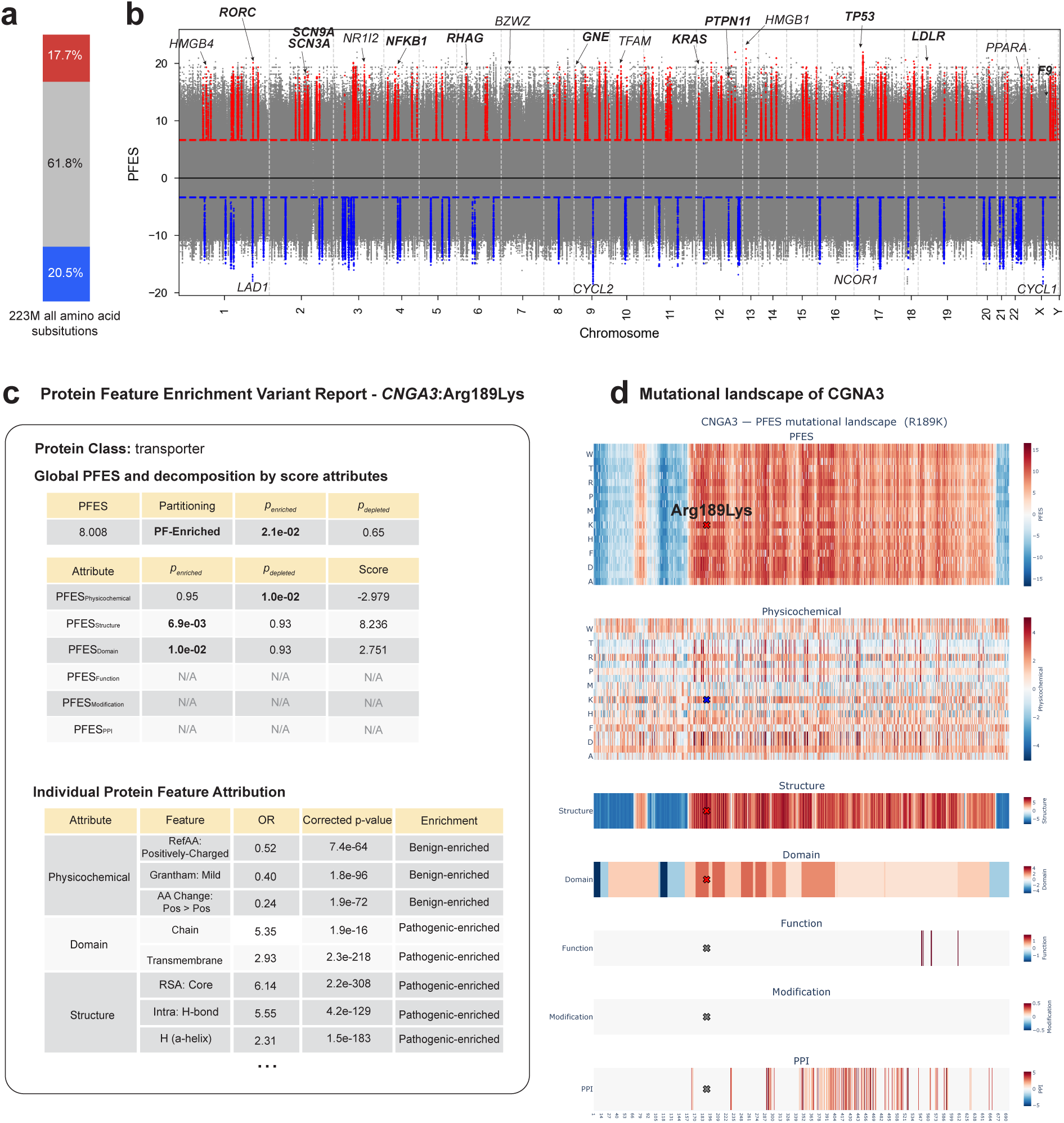
PFES as a proteome-wide variant interpretation resource. **(a)** Proteome-wide partitioning of all 223 million possible amino acid substitutions from 20,132 human proteins. Fractions of PF-Enriched (red, 17.7%), PF-Neutral (gray, 61.8%), and PF-Depleted (blue, 20.5%) variants are shown. **(b)** Residue-level protein feature enrichment landscape plotted by chromosomal position. Each point represents the median PFES across 19 different substitutions at a given residue position (11,396,719 amino acid positions in total). Prioritized genes are highlighted in red (125 PF-Enriched genes) or blue (87 PF-Depleted genes). Dashed lines indicate the PFES thresholds for partitioning: PFES > 6.65 (PF-Enriched, red) and PFES < −3.36 (PF- Depleted, blue). Selected gene names are labeled as examples, with bold font indicating genes with strong or definitive gene-disease relationships in GenCC and normal font indicating genes not currently represented in GenCC. **(c)** The example of protein feature enrichment variant report is shown for *CNGA3* p.Arg189Lys, generated from Google Colab.The report displays the global PFES, its partitioning category with associated statistical significance (*p*_enriched_ and *p*_depleted_), and subscore attributes from the six protein feature groups. A table below shows individual protein features associated with this variant, along with its odds ratios (OR), corrected *p*-value, and its enrichment direction. **(d)** Protein-level mutational landscape heatmap for *CNGA3*. The top panel shows the global PFES landscape for every possible amino acid substitution, with the queried variant Arg189Lys marked. Panels below decompose the landscape by score attributes, revealing positional heterogeneity in protein feature enrichment across the protein sequence. Positive scores (red) indicate enrichment for pathogenic-associated protein features; negative scores (blue) indicate enrichment for tolerance-associated features.

To examine how protein feature enrichment varies at the gene level, we computed the median PFES across all possible substitutions at each residue positions (11.4 million) and plotted these by chromosomal location (**Fig. 7b**). Genes whose fraction of PF-Enriched or PF-Depleted substitutions fell within the top 5% of their chromosome and whose median PFES reached statistical significance against the corresponding reference distribution (*p* < 0.0001) were identified as genes whose mutational landscape is dominated by either pathogenic- or tolerance-associated protein feature profiles. This identified 125 PF-Enriched and 87 PF-Depleted genes (**Supplementary Table 4**). PF-Enriched genes were significantly overrepresented among established rare disease genes in the Gene Curation Coalition (GenCC) database (*62*) (OR = 3.04, p = 5.34e-9; **Fig. 7b**, *boldfaced*), while PF-Depleted genes showed no significant enrichment. A subset of the 125 PF-Enriched genes is not yet represented in GenCC (e.g., *NR1I2;* nuclear receptor, *TFAM*; mitochondrial transcription factor), representing candidates whose protein feature architecture suggests functional constraint not yet captured by existing disease gene catalogs.

To enable community use, we have published a precomputed PFES for 223M all possible missense variants across 20,245 human proteins, as well as a readily available resource through a Google Colab notebook where users can query a missense variant of interest and retrieve the complete output (**Fig. 7c-d; Data Availability**). The notebook retrieves protein annotations from the G2P portal (*42*) at query time, ensuring access to current annotations. For any queried variant, two outputs are generated. The first is a per-variant PFES report (**Fig. 7c**), illustrated for *CNGA3*:Arg189Lys: global PFES (8.0), partitioning category (PF-Enriched, *p*_enriched_ = 2.1e-2), and subscores from all six attribute groups. Structural features (PFES_Structure_ = 8.236, p = 6.9×10⁻³) and domain annotations (PFES_Domain_ = 2.751, p = 1.0e-2) drive the enrichment, while the physicochemical subscore is depleted (PFES_Physicochemical_ = −2.979, p = 1.0e-2), reflecting the charge-preserving nature of the substitution. An individual feature attribution table lists each contributing feature with its odds ratio and FDR-corrected p-value. The second output is a protein-wide mutational landscape heatmap (**Fig. 7d**): the top panel shows PFES across all possible substitutions at all positions, contextualizing the queried variant within the full substitution space, and the six panels below decompose this landscape by attribute group. Together, these outputs allow users to interpret a variant in terms of which protein characteristics are implicated and how its profile compares to the broader mutational landscape of the protein.

## Discussion

Technological advances have made genome and exome sequencing as well as the identification of variants accessible to patients (*63*). While computational variant effect predictors (VEPs) can classify these variants into pathogenic or benign, they often fail to answer the demanding question of what molecular context makes a variant pathogenic (*64*). The Protein Feature Enrichment Score (PFES) addresses this need directly by quantifying the molecular context of missense variants in terms of their protein features across 20 protein functional classes and providing decomposition of the overall PFES score into interpretable feature attributions alongside classification (**Fig. 1**).

Prior work by Iqbal *et al.* (*18*) established that enrichment of 3D protein structural features distinguishes pathogenic from population variants, but using experimental structures only, limiting both proteome coverage and statistical power needed for optimization. By leveraging AlphaFold2 structural models, we were able to optimize the PFES framework, which demonstrated that not all variants in gnomAD, but only common variants (allele frequency > 1%) combined with ClinVar benign variants, provide a stringent baseline for identifying comparatively pathogenic-enriched protein features (**Fig 2**). The proteome-wide enrichment analysis also revealed previously uncharacterized insights about the degree of class-specificity in pathogenic variant profiles (**Fig. 3b**). Enzymatic and transport-related classes, such as metabolite interconversion enzyme, DNA metabolism protein, and transporter, show pathogenic profiles close to the proteome-wide baseline, as they tend to operate through catalytic or substrate-handling mechanisms that are broadly shared. In contrast, classes with high deviation, including extracellular matrix protein, structural protein, and cell adhesion molecule, are predominantly involved in tissue architecture or macromolecular assembly, where functional requirements are more specialized and less transferable across protein families. At the feature level, physicochemical severity is enriched among pathogenic variants across all classes: a universal constraint on amino acid compatibility that operates independently of functional context. Other features are meaningful only where they are functionally relevant: DNA-binding region enrichment is confined to transcriptional regulators and DNA metabolism proteins (**Fig. 3d**). Gerasimavicius *et al.* (*60*) arrived at a consistent conclusion from a complementary direction: the prevalence of stabilizing and destabilizing variants in haploinsufficiency disease genes varies systematically across different protein classes. Altogether, these reported results and ours demonstrate that the pathogenic relevance of individual protein features is not fixed but shaped by the functional roles of proteins.

The performance of PFES relative to existing predictors is best understood in the context of what the framework is designed to measure. Benchmarking against deep mutational scanning (DMS) datasets, which have become the primary standard for comparative evaluation in the field, showed that PFES achieves a moderate average DMS correlation (ρ = 0.368) compared to the top performers including AlphaMissense (ρ = 0.525), SaProt (ρ = 0.520), VARITY_R (ρ = 0.512), and CPT-1 (ρ = 0.510) (**Supplementary Table 2**; **Fig. 4c**). This performance gap reflects two distinct sources. The first is a deliberate trade-off: PFES derives scores from explicit protein feature annotations rather than optimization against labeled training data, prioritizing interpretability over classification accuracy. The second is a position-level resolution ceiling: because PFES values are primarily tied to protein features at the reference amino acid position, substitutions at the same position receive similar values regardless of the specific amino acid change introduced. This limitation is not unique to PFES; it has been observed across current variant effect predictors broadly that VEPs consistently assign a much narrower range of scores to substitutions at a given reference amino acid position than DMS experiments do. It reflects an unresolved challenge in distinguishing the specific amino acid change from the position being changed (*65*). PFES is therefore better suited to characterizing the protein feature sensitivity landscape across positions within a protein than to distinguish between individual substitutions at a single site, consistent with its stronger agreement with full-length DMS datasets (**Fig. 4c**; **Supplementary Fig. 7**). Performance heterogeneity across genes is also a shared characteristic of all predictors, with no single tool performing uniformly well across all proteins. For PFES specifically, gene-level variation in DMS correlation carries a distinct interpretive meaning beyond performance differences: strong agreement indicates that a protein’s functional sensitivity is well-captured by protein annotations, whereas poor agreement points towards mechanisms operating outside the scope of currently annotated protein features, which itself warrants further investigation rather than simply reflecting a prediction failure.

In VUS triage, PFES is most useful when considered alongside existing variant effect predictors rather than in isolation. A variant that receives a high pathogenicity probability from AlphaMissense or REVEL and is additionally PF-Enriched carries a more interpretable evidence profile: the former answers whether the variant is likely pathogenic, while PFES explains which protein characteristics are implicated (**Figs. 5c-e**). Conversely, a variant scored as pathogenic by existing tools but falling in the PF-Neutral category may prompt closer examination of whether its mechanism involves features outside current annotation coverage. Importantly, PF-Neutral status should not be interpreted as evidence of benignity; it reflects only that the variant lacks a distinctive protein feature profile in either direction. Where existing predictors provide a ranked list of variants according to the score, PFES decomposition can help investigators understand the protein-level basis for that ranking and design functional assays accordingly (**Fig. 1d**, **Fig 4a-b**). The utility of attribute-level decomposition extends to mechanistic distinction between loss-of-function (LoF) and gain-of-function (GoF) variants (**Fig. 6**). Gerasimavicius *et al.* (*60*) showed that GoF and dominant-negative mutations have profoundly different structural profiles from LoF mutations: milder effects on overall protein stability but strong enrichment at protein-protein interaction interfaces. PFES recapitulates this pattern across 596 genes from Gene2Phenotype (*61*): PPI features were nearly three times more prevalent in GoF variants (∼18%) than in LoF variants (∼6.5%), while overall PFES distributions were broadly overlapping, confirming that the difference is mechanistic rather than a general pathogenicity signal (**Fig. 6b-c**). This is consistent with GoF mechanisms operating more frequently through aberrant complex formation or interface perturbation than through protein destabilization(*59, 60*).

The recognition that variant classification benefits from mechanistic interpretation has motivated a growing number of proteome-wide approaches. MutPred2(*55*) established an early foundation by modeling the impact of amino acid substitutions on over 50 structural and functional protein properties. Jänes *et al*.(*66*) computed stability changes for over 200 million variants and characterized protein-protein interface and binding pocket disruption using AlphaFold2-predicted structures and complexes. Cagiada *et al*.(*57*) addressed the distinction between two principal routes to loss-of-function variants: whether pathogenicity arises through protein destabilization or direct disruption of functional interactions. PFES does not make this stability-versus-function distinction explicitly, but its structural and functional attribute decomposition provides complementary information: whether a variant occurs at a structurally constrained position, a functional site, a post-translational modification site, or a protein-protein interaction interface, each assessed against the enrichment background of the protein’s functional class (**Fig. 2f**; **Fig. 3a**).

Beyond variant prioritization, PFES supports gene-level prioritization for rare disease through a direction distinct from sequence-based constraint (*67*), residue-level missense constraint (*68*), and phenotype-driven candidate ranking (*69*), all of which are agnostic to protein-level disease mechanism. Genes with a high proportion of PF-Enriched variants are more likely to cause disease through disruption of specific structural or functional protein characteristics, a signal that is most interpretable in rare disease genes where strong selective constraint means single missense variants are sufficient to cause disease. Consistent with this, PF-Enriched genes are significantly overrepresented among well-established rare disease genes in GenCC (**Fig. 7b**), establishing protein feature enrichment as a gene-level evidence layer orthogonal to existing prioritization approaches.

PFES is inherently dependent on existing protein annotations from PDB, AlphaFoldDB, UniProtKB, PhosphoSitePlus, and PANTHER. Proteins that are poorly characterized structurally or functionally will yield lower-resolution profiles, and unannotated functional sites, novel interaction interfaces, or PTMs absent from current databases will not contribute to the score. Variants acting through highly protein-specific mechanisms, subtle allosteric perturbations not captured in current annotations, or gain-of-function effects involving novel interactions will often fall in the PF-Neutral category because their pathogenic mechanism lies outside the scope of features currently annotated. Finally, enrichment patterns are derived from ClinVar, HGMD and gnomAD, datasets that are not uniformly distributed across the proteome and reflect the diseases and genes for which genetic diagnosis is most actively pursued. Protein classes underrepresented in clinical databases may have less robust enrichment estimates.

The annotation-dependence that defines PFES’s current boundaries is also the source of its extensibility. Unlike models that require retraining on new labeled data, PFES improves in coverage and resolution as annotations expand: new feature categories can be incorporated directly into the enrichment analysis without architectural changes to the framework. This extensibility operates not only through passive growth of proteome-wide databases but through deliberate, targeted application of the same enrichment logic to specific disease contexts or gene sets. Applying protein feature enrichment analysis to curated structural resources for neurodevelopmental disorder genes (*43*), for instance, identified functionally essential 3D sites with 8-fold enrichment of patient variants, while directing the same approach to ALS-associated genes (*70*) revealed structural and physicochemical enrichment patterns distinct from those seen in neurodevelopmental disorders. These findings demonstrate that disease-specific instantiation produces biologically distinct insights rather than simply recapitulating proteome-wide findings. As protein annotation databases continue to grow through structural genomics, proteomics, and high-throughput functional studies, the scope of mechanisms detectable by PFES will broaden accordingly.

## Methods

### Missense variant collection from ClinVar, HGMD, and gnomAD

Missense variants annotated on canonical protein isoforms were collected from three primary sources: ClinVar, Human Gene Mutation, and the Genome Aggregation Database (gnomAD). Variant data from ClinVar was downloaded directly from the FTP site (https://ftp.ncbi.nlm.nih.gov/pub/clinvar/tab_delimited/variant_summary.txt.gz). Single nucleotide variants resulting in missense changes on the reference genome GRCh38 were collected. From the Human Gene Mutation Database (HGMD®) professional release (version 2025.1), single nucleotide variants resulting in missense changes on GRCh38 and with disease-causing state (variantType = ‘DM’ flag, indicating disease mutation) were extracted. For gnomAD variant collection, we downloaded raw VCF files (https://gnomad.broadinstitute.org/downloads/) for genome and exome datasets from gnomAD v4.1.1 and selectively extracted variants that passed all variant filters for quality control (filter=’PASS’ flag). Then, single nucleotide variants resulting in missense changes were collected. When the same variant was identified in both genome and exome datasets, we summed the allele count and the allele number, subsequently calculating the merged allele frequency. We defined four missense variant datasets for statistical analysis as follows (see **Fig. 1a**).

(1) Pathogenic: ClinVar variants annotated as pathogenic or likely pathogenic (P/LP) and HGMD disease variants annotated as high confidence, indicating variants with proven disease-causing roles, supported by functional evidence in the literature. To ensure dataset independence, variants also present in gnomAD were excluded (*n*=51,919 removed). This exclusion does not imply these variants are non-pathogenic, but may represent variants in autosomal recessive conditions or of incomplete penetrance. After exclusion, 85,321 pathogenic missense variants were retained.
(2) Benign: ClinVar variants annotated as benign or likely benign (B/LB). Variants present also in gnomAD were excluded, as high allele frequency in population database already constitutes a criterion for benign classification under ACMG/AMP guidelines (BA1, BS1), and retaining them would introduce redundancy and overlap with population datasets. This ensures retained benign variants are supported by clinical evidence beyond population frequency alone. After exclusion, 104,848 benign missense variants were retained.
(3) Population: gnomAD variants, further divided into allele frequency (AF) subgroups: Rare (AF < 0.1%), Intermediate (0.1% ≤ AF < 1%), and Common (AF ≥ 1%). In total, 12,089,818 missense variants were collected. Variants overlapping with the pathogenic dataset were excluded as described above.
(4) VUS: ClinVar variants of uncertain significance, totaling 1,489,214.

Because our analysis focuses on protein-level effects, each unique amino acid substitution (e.g., His123Gln in a given protein) is counted once regardless of how many distinct nucleotide changes produce it at the genomic level.

### Protein feature annotations

Each variant position was annotated with 103 protein features spanning six attribute categories: physicochemical properties, 3D structural features, domain and region annotations, functional site annotations, post-translational modifications, and protein-protein interaction residues. Features were collected from public databases or precomputed as described below.

#### Physicochemical properties (count: 46)

Each amino acid substitution was characterized by three physicochemical descriptors. First, the reference residue was assigned to one of seven side chain classes: *Aliphatic* (Ala, Ile, Leu, Met, Val), *Aromatic* (Phe, Trp, Tyr), *Polar/Neutral* (Asn, Gln, Ser, Thr), *Positively-Charged* (Arg, His, Lys), *Negatively-Charged* (Asp, Glu), and *Special* (Pro, Gly, Cys). Second, the substitution type was recorded as the residue class transition (e.g., *Polar/Neutral*>*Negatively-Charged*). Third, the severity of the physicochemical shift was quantified using Grantham’s distance, which considers amino acid composition, polarity, and molecular volume, and classified into four categories: *Mild* (D < 50), *Moderate* (50 ≤ D < 100), *Substantial* (100 ≤ D < 150), and *Severe* (D ≥ 150).

#### 3D structural features (count: 22)

Structural features were computed from AlphaFold2 predicted structures and available PDB structures. Secondary structure was assigned using the DSSP algorithm applied to AlphaFold2 structures, yielding nine categories: *Alpha-helix (H), 3_10_-helix (G), Polyproline helix (P), π-helix (I), Isolated beta-bridge (B), Extended beta-sheet (E), Turn (T), Bend (S),* and *Coil (C).* Relative solvent accessibility (RSA) was calculated as the DSSP-derived absolute solvent accessible surface area normalized by the maximum possible exposure of each residue in a Gly-X-Gly reference tripeptide configuration, using reference values (*71*). RSA values were categorized into five exposure levels: *Core* (RSA < 5%), *Buried* (5–25%), *Medium-buried* (25–50%), *Medium-exposed* (50–75%), and *Exposed* (≥ 75%). Structural confidence was characterized using the AlphaFold2 per-residue pLDDT score, classified as *Very high* (> 90), *Confident* (70–90), *Low* (50–70), or *Very low* (≤ 50). Intramolecular interactions were detected from AlphaFold2 and monomeric PDB structures using HBPLUS with the following distance-based criteria: *Intramolecular hydrogen bonds* were identified when the donor-acceptor distance was less than 3.9 Å and the hydrogen-acceptor distance was less than 2.5 Å; *Intramolecular VdW interactions* (van der Waals contacts) were identified when the distance between two residues was less than 3.9 Å;

*Intramolecular disulfide bonds* were identified when the distance between sulfur atoms of two cysteine residues was less than 3.0 Å; and *Intramolecular salt bridges* were identified when the distance between oppositely charged atoms of Asp, Glu, Arg, or Lys residues was less than 3.2 Å. For AlphaFold2-derived interactions, an additional Predicted Aligned Error (PAE) filter of 8 Å was applied to exclude contacts between residues with low positional confidence. Intramolecular disulfide bonds additionally include UniProtKB disulfide bond annotations classified as intramolecular.

#### Functional site annotations (count: 5)

Functional site annotations were retrieved from UniProtKB and include *Active site*, *Binding site*, *DNA-binding region*, *Zinc finger*, and *Site*. The last covering single-residue annotations of functional interest such as cleavage or inhibitory sites.

#### Domain and region annotations (count: 16)

Domain and region annotations were retrieved from UniProtKB and include *Domain*, *Transmembrane*, *Intramembrane*, *Topological domain*, *Signal peptide*, *Transit peptide*, *Propeptide*, *Coiled coil*, *Repeat*, *Motif*, *Peptide*, and *Chain*. UniProtKB Region annotations were further subdivided into three categories: *Region/Disordered* (intrinsically disordered regions), *Region/Interaction* (regions annotated for intermolecular interactions), and *Region/Others* (miscellaneous annotations).

#### Post-translational modifications (count: 10)

PTM annotations were collected from two sources. From UniProtKB, *Lipidation*, *Glycosylation*, *Crosslinks*, and *Modified residue* were retrieved. From PhosphoSitePlus, site-level annotations for *Acetylation*, *Methylation*, *Phosphorylation*, *SUMOylation*, *Ubiquitination*, and *O-GalNAc/GlcNAc* glycosylation were collected.

#### Protein-protein interaction features (count: 4)

Protein-protein interactions (PPI) were detected from experimentally resolved protein complex structures in the PDB. The same distance-based criteria is used to define *Intermolecular hydrogen bond*, *Intermolecular salt bridge*, *Intermolecular disulfide bond*, and *Intermolecular VdW interaction* but exclusively between residues in different proteins. *Intermolecular disulfide bonds* additionally include UniProtKB disulfide bond annotations tagged as interchain. t

### Fisher’s exact test and Benjamini-Hochberg procedure

Two-sided Fisher’s exact tests were performed for each binary or categorical feature within each PANTHER protein functional class. For each feature, a 2×2 contingency table was constructed comparing the presence or absence of that feature in pathogenic variants versus control variants. Odds ratios (OR) with 95% confidence intervals were calculated from the contingency table entries. P-values from Fisher’s exact tests were corrected for multiple comparisons within each protein class using the Benjamini-Hochberg false discovery rate (FDR) procedure. Features with FDR-corrected *p*-values, i.e. *q*-values, below 0.01 were considered significantly enriched. The stricter threshold of 0.01 was chosen to reduce the risk of false positive enrichment signals, given the large number of features tested simultaneously across protein classes. Features that did not meet the FDR threshold were excluded from the PFES calculation and contributed no log odds ratio to the score.

### Computation of Protein Feature Enrichment Score (PFES)

The Protein Feature Enrichment Score (PFES) is a quantitative measure of protein feature signature enrichment associated with pathogenic or benign variation, enabling mechanistic interpretation of missense variants. For a given variant, PFES is calculated as the sum of log odds ratios across all significantly enriched features: 𝑃𝐹𝐸𝑆 = 𝛴 𝑙𝑜𝑔(𝑂𝑅_*i*_), where the summation is taken over all features *i* that are significantly enriched (FDR-corrected *p* < 0.01) in either the pathogenic or control group. Features enriched among pathogenic variants contribute positive log OR values; features enriched among control variants contribute negative log OR values. Consequently, a positive PFES indicates that the variant’s protein characteristic profile is enriched for features associated with pathogenic variants, while a negative PFES indicates enrichment for features associated with benign variants. When an OR is zero or undefined due to complete depletion or enrichment of a feature, it is replaced by the minimum or maximum finite OR observed across all features within the given protein class, respectively. This ensures that features with the strongest effect sizes are not excluded from the score due to numerical instability.

PFES is further decomposed into six attribute-level sub-scores (PFES_Physicochemical_, PFES_Structure_, PFES_Domain_, PFES_Function_, PFES_Modification_, and PFES_PPI_) by summing the log OR contributions within each feature category separately. This decomposition supports mechanistic interpretation by identifying which categories of protein characteristics drive the overall enrichment signal for a given variant.

### PFES-based partitioning of PF-Enriched, Neutral, and Depleted variants

Variants were partitioned into three categories based on the statistical deviation of their PFES from the empirical distributions of clinically annotated pathogenic and control variants. Empirical PFES distributions were constructed separately for pathogenic (case, n = 85,321) and benign and common population (control, n = 130,719) variants and smoothed using kernel density estimation (KDE).

Two one-sided p-values were derived for each variant from the tail areas of the KDE distributions at the observed PFES. *p_enriched_* is the right-tail area under the control KDE at the variant’s PFES, quantifying how enriched the variant’s protein feature profile is relative to control variation – a small value indicates the score falls in the upper tail of the control distribution, where pathogenic-associated protein features are overrepresented. *p_depleted_* is the left-tail area under the case KDE at the variant’s PFES, quantifying how depleted the variant’s protein feature profile is relative to pathogenic variation –a small value indicates the score falls in the lower tail of the case distribution, where tolerance-associated protein features are overrepresented. A variant was classified as *PF-Enriched* if *p_enriched_* < 0.05, *PF-Depleted* if *p_depleted_* < 0.05, and *PF-Neutral* if both p-values were ≥ 0.05, reflecting the absence of a distinctive protein feature signal in either direction.

The same partitioning logic applies at the individual attribute level, where each attribute sub-score is independently compared against its own empirical case and control distributions, yielding attribute-specific enrichment categories (see **Supplementary Fig. 4**). The derived partitioning thresholds are as follows: PFES > 6.664 (PF-Enriched) and PFES < −3.351 (PF-Depleted); PFES_Physicochemical_ > 2.169 (Enriched) and < −2.280 (Depleted); PFES_Structure_ > 6.353 (Enriched) and < −2.785 (Depleted); PFES_Domain_ > 1.842 (Enriched) and < −2.556 (Depleted); PFES_Modification_ > 0.009 (Enriched) and < −0.042 (Depleted); PFES_Function_ > 0.0 (Enriched only); PFES_PPI_ > 0.0 (Enriched only).

### Pairwise feature distribution comparison: Kolmogorov-Smirnov test and Cohen’s w

To quantify distributional differences in protein features between variant groups, two complementary statistics were used. For continuous features (e.g., RSA, pLDDT, Grantham distance), the Kolmogorov-Smirnov (KS) test was applied to each pairwise combination of variant groups. The KS statistic reflects the maximum absolute difference between the empirical cumulative distribution functions of the two groups, ranging from 0 (identical distributions) to 1 (completely non-overlapping distributions).

For binary and categorical features, pairwise distributional divergence was quantified using a normalized Cohen’s w. For each feature pair, Cohen’s w was computed from the proportion distributions of the two variant groups as 𝑤 = √(𝛴(𝑝₁ᵢ − 𝑝₂ᵢ)² / 𝑟ᵢ) / √(𝑘 − 1), where p₁ᵢ and p₂ᵢ are the observed proportions of category *i* in each group, *rᵢ* is the mean proportion across both groups serving as the reference distribution, and k is the number of categories. Dividing by √(𝑘 − 1) bounds the statistic to [0, 1] regardless of category count where 0 indicates identical distributions, and 1 indicates complete non-overlap, enabling direct comparison across features with different numbers of categories., enabling direct comparison across features of varying complexity, for example, a binary feature (k = 2, normalization factor = 1) versus secondary structure (k = 9, normalization factor = √8 ≈ 2.83). Mean KS and Cohen’s w values across all features were used to assess overall protein feature divergence between datasets and inform control group selection.

### Protein class annotation from PANTHER

Each protein in the dataset was assigned to a functional class using the PANTHER (*38*) (Protein ANalysis THrough Evolutionary Relationships; https://pantherdb.org/) classification system (version 19.0), which defines 24 protein classes at the highest level of its hierarchical taxonomy. Variants were mapped to their corresponding protein class via UniProt accession identifiers. Proteins that could not be assigned to any PANTHER class were retained as an *unclassified* category (**Supplementary Fig. 3**) . For statistical power, protein classes with fewer than 300 variants in either the pathogenic or control dataset were merged into this unclassified category. Five classes met this criterion: cell junction protein, chaperone, calcium-binding protein, viral or transposable element protein, and storage protein. In total, 20 protein classes, including the consolidated unclassified group, were retained for class-stratified analyses.

### Benchmarking with deep mutational scanning data and variant effect predictors

To benchmark PFES against established variant effect predictors, we used the curated DMS dataset compiled by Livesey and Marsh (*51*), which provides VEP predictions across 36 proteins with deep mutational scanning data. PFES was compared against 13 variant effect predictors: AlphaMissense, CPT, VARITY_R, REVEL, SIFT, PolyPhen-2 (HumVar), CADD (v1.7), SIGMA, AlphScore, Rhapsody, SNPs&GO3D, SaProt, and MutPred. For each DMS dataset, Spearman’s rank correlation coefficient (ρ) was calculated between each predictor’s scores and the experimental fitness scores across all covered variants. For proteins with multiple independent DMS datasets, a single score set was chosen as the set with the highest median Spearman’s correlation across all 13 predictors, preventing outliers from unduly influencing dataset selection.

### Evaluation of statistical concordance

For variant classification concordance analyses comparing PFES partitioning against VUS subtiers provided by the Mass General Brigham Laboratory for Molecular Medicine, statistical agreement was quantified using Kendall’s rank correlation coefficient (τ). Kendall’s τ measures the association between two ordinal rankings by comparing the proportion of concordant and discordant pairs, and is appropriate for ordered categorical data with potential ties. For each gene with sufficient variant representation, τ was calculated between PFES partition category (PF-Enriched, PF-Neutral, PF-Depleted, encoded as ordinal ranks) and the VUS subtiers (VUS-high, VUS-mid, VUS-low, encoded as ordinal ranks). Genes were included in the analysis if they contained at least 2 variants in each of the VUS-high and VUS-low tiers, ensuring sufficient representation at both ends of the pathogenicity spectrum for a meaningful concordance assessment. p-values were derived from the normal approximation to the null distribution of τ under the assumption of no true association, with tie correction applied.

### Collection of Loss- and Gain-of-function variants from Gene2Phenotype

Loss-of-function (LoF) and gain-of-function (GoF) variant datasets were derived from Gene2Phenotype (*61*) (https://www.ebi.ac.uk/gene2phenotype), a curated database of gene-disease associations annotated with disease mechanism. Gene-disease records were retrieved from the Developmental Disorders panel, which represents the largest and most comprehensively annotated panel. Records were filtered to retain only those with molecular mechanism annotated as ‘loss of function’ or ‘gain of function’, variant consequence annotated as ‘altered gene product structure’. This yielded 150 LoF and 158 GoF gene-disease records from 2,822 total records. For each retained gene-disease record, corresponding pathogenic missense variants were retrieved from ClinVar (P/LP) and HGMD (disease mutation, high confidence), and matched to the annotated disease if available. *FBN1* was excluded from the LoF dataset due to the absence of a complete AlphaFold2 structure (2,821 amino acids) and because its variants alone comprised more than 50% of the entire LoF dataset, which would have introduced substantial bias. After phenotype matching and exclusions, 1,284 LoF and 1,753 GoF pathogenic missense variants were retained among 82 and 117 genes, respectively. A control group was constructed from our control dataset (benign and common population variants) mapped to the same genes, resulting in 4,097 variants among 176 genes. This dataset was used to evaluate whether PFES captures protein feature signatures specific to disease mechanism, and to assess how feature category composition differs between LoF and GoF variant groups.

## Acknowledgement

We thank Patrick May (Luxembourg Centre for Systems Biomedicine, University of Luxembourg) for providing the HGMD professional data (version 2025.1).

## Author contributions

S.I. designed the study and supervised the project. S.K. led the project, performed the analysis, and developed the code and dataset. J.S. performed data curation from the G2P portal. S.I and S.K. wrote the manuscript. M.D., M.L., and H.L.R provided data, discussion, and reviewed the manuscript. S.I. contributed to the funding acquisition.

## Data availability

Protein feature and genetic variant data from ClinVar and gnomAD are available to retrieve from the Genomics 2 Proteins portal website (https://g2p.broadinstitute.org/). Proteome-wide precomputed PFES is made available from Zenodo (https://zenodo.org/records/19710599). Colab notebook for PFES computation (https://colab.research.google.com/github/broadinstitute/missense-pfes/blob/main/notebooks/G2P_PFES.ipynb). Codes for data analysis and Python client library are available in the GitHub repository (https://github.com/broadinstitute/missense-pfes#).

