## Supplementary Figures 1-8 for "Human Proteome-wide Mechanistic Interpretation of Missense Variants through Protein Feature Enrichment Score"

**Supplementary Tables and Figures**

**Supplementary Table 1.** Protein class-specific odds ratios (OR), p-values, and corrected p-values for the entire protein features across protein classes.

**Supplementary Table 2.** Spearman correlation coefficients between PFES and 13 variant effect predictors against 36 deep mutational scanning datasets.

**Supplementary Table 3.** Gene-Disease relationship by molecular mechanism (loss- or gain-of-function) curated from Gene2Phenotype database.

**Supplementary Table 4**. Gene-level summary of PFES in the human proteome.

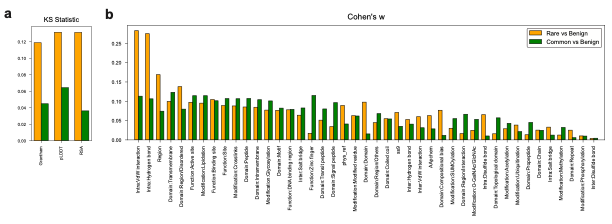

**Supplementary Fig. 1. Per-feature distributional divergence between benign and rare versus benign and common population variants. (a)** KS statistics for continuous features (AA distance, pLDDT, and RSA) comparing rare versus benign (orange) and common versus benign (green) variant pairs. For all three features, divergence is smaller between common and benign than between rare and benign variants. **(b)** Normalized Cohen's w for all binary and categorical features comparing rare vs. benign (orange) and common vs. benign (green) variant pairs. For the majority of features, divergence is smaller between common and benign variants, supporting common population variants as the most appropriate proxy-benign group.

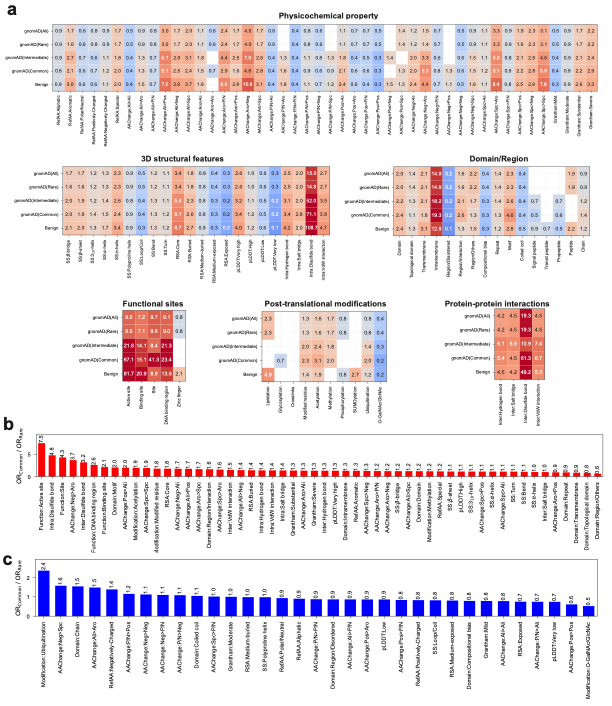
**Supplementary Fig. 2. Odds ratios of protein features in pathogenic variants relative to each gnomAD allele frequency spectrum. (a)** Heatmaps of OR values for pathogenic variants relative to different control groups: gnomAD(All), gnomAD(Rare), gnomAD(Intermediate), gnomAD(Common), and ClinVar benign across all six protein feature categories – physicochemical properties, 3D structural features, domain/region, functional sites, post-translational modifications, and protein-protein interactions. Red indicates enrichment in pathogenic variants (OR > 1) and blue indicates depletion (OR < 1). White cells indicate insufficient data for OR estimation. **(b)** Ratio of OR using common population variants as control to OR using all gnomAD variants as control (OR_Common_ / OR_All_) for features enriched in pathogenic variants, sorted in descending order. Values above 1 indicate that the signal is amplified when common variants are used as the control group. Functionally constrained features such as active sites, disulfide bonds, and functional sites show the greatest amplification. **(c)** Ratio of OR_Common_ / OR_All_ for features depleted in pathogenic variants, sorted in descending order. Values below 1 indicate further depletion when common variants serve as the control, with O-GalNAc/GlcNAc and exposed positions showing the greatest change.

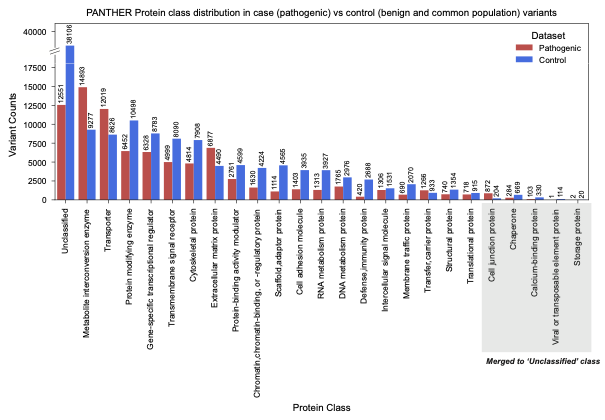

**Supplementary Fig. 3. PANTHER functional protein classes and variant counts by protein class.** The number of pathogenic (red) and control, i.e., benign and common (blue) variants, is shown for each PANTHER protein class used for the enrichment analysis. Five protein classes with fewer than 300 variants in either the pathogenic or control dataset (i.e., cell junction protein, chaperone, calcium-binding protein, viral or transposable element protein, and storage protein) were merged into the unclassified class for all subsequent analyses.

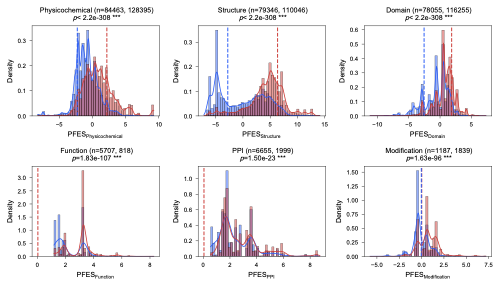

**Supplementary Fig. 4. Distribution of decomposed PFES from pathogenic and control variants.** Density distributions of subscores from each of the six attribute categories: physicochemical, structure, domain, function, PPI, and modification, comparing pathogenic (red) and control (blue) variants. Sample sizes (*n*) for pathogenic and control variants are indicated for each attribute. Dashed lines indicate the score at which one-sided *p*-value (*p*_enriched_ and *p*_depleted_) becomes less than 0.05. For functional and PPI attributes, presence alone serves as the enrichment criterion given their unidirectional score distributions. Statistical significance of distribution differences assessed by the Mann-Whitney U test; *p*-values are shown.

**Supplementary Fig. 5. Residue-level decomposed PFES attribute profiles for *PPARG*.** Score profiles along the *PPARG*/P37231 protein sequence (*n*=9,595 variants) for DMS functional readouts and each PFES attribute: physicochemical, structure, function, domain, PPI, and modification. Points are colored by score magnitude. Several annotated features contributing to functional, domain, and modification attribute scores are labeled, including DNA-binding region, binding sites, region/interaction, lipid-binding domain, and ubiquitination sites.

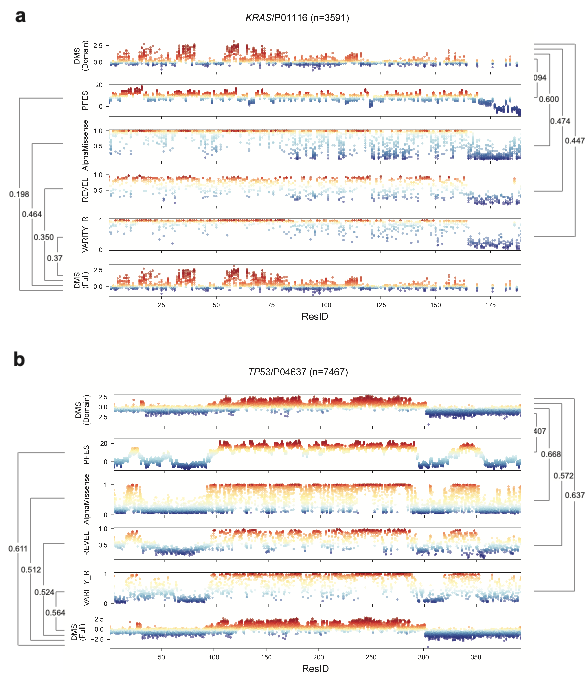

**Supplementary Fig. 6. Effect of DMS dataset coverage on benchmarking performance for *KRAS* and *TP53*.** Residue-level score profiles comparing domain-focused and full-length DMS datasets against PFES, and variant effect predictors (VEPs) – AlphaMissense, VARITY_R, and REVEL for **(a)** *KRAS*/P01116 (n=3,591) and **(b)** *TP53*/P04637 (n=7,467). Top and bottom tracks are shown for two DMS tracks: the domain-focused dataset selected by standard protocol (DMS (Domain)) and the full-length dataset covering the entire protein sequence (DMS (Full)), respectively. Points are colored by score magnitude. Spearman's correlation values between each score and the respective DMS dataset are shown. PFES is the only score showing increased correlation when compared with the full-length DMS dataset rather than with the domain-focused DMS dataset.

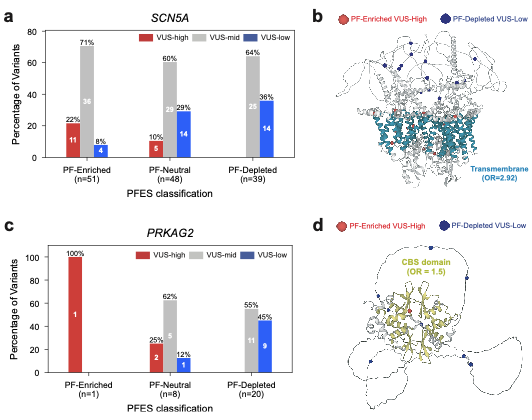

**Supplementary Fig. 7. Gene-level concordance between PFES partitioning and VUS subtier classification for *SCN5A* and *PRKAG2*. (a)** Distribution of VUS subtiers (VUS-high, VUS-mid, VUS-low) across PFES partition categories (PF-Enriched, PF-Neutral, PF-Depleted) for *SCN5A* (*n*=138). 22% of PF-Enriched variants overlap with VUS-high, while PF-Depleted variants include no VUS-high. The fraction of VUS-low variants is highest among PF-Depleted variants (36%), decreasing for PF-Neutral (29%), and PF-Enriched (8%) variants. **(b)** Structural mapping shows PF-Enriched VUS-high variants (red) clustering within the transmembrane domain (teal; OR = 2.92), and PF-Depleted VUS-low variants (blue) distributed in disordered regions outside of the transmembrane helical bundle. **(c)** Distribution of VUS subtiers across PFES partition categories for *PRKAG2* (*n*=29). The single PF-Enriched VUS is classified as VUS-high (100%), while PF-Depleted variants are predominantly VUS-low (45%). **(d)** Structural mapping (right) shows PF-Enriched VUS-High variants (red) clustering within the cystathionine-beta-synthase (CBS) domain (yellow).

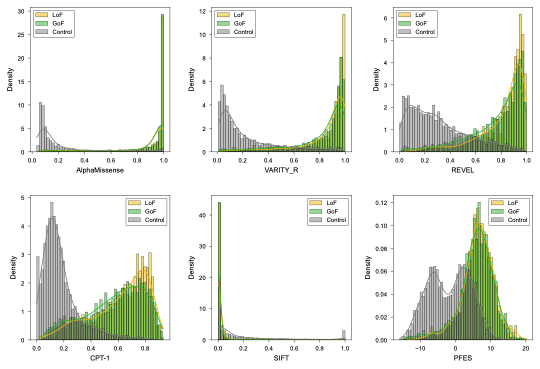
**Supplementary Fig. 8. Variant effect predictor (VEP) scores do not distinguish loss-of-function from gain-of-function variants.** Score distributions of five variant effect predictors – AlphaMissense, VARITY_R, REVEL, CPT-1, SIFT – and PFES are shown separately for loss-of-function (LoF, yellow) and gain-of-function (GoF, green) variants. For all six scores, LoF and GoF distributions are broadly overlapping, indicating that none of these tools captures the disease mechanism by a score alone. PFES distributions for LoF and GoF variants are also nearly identical, centered around positive values consistent with pathogenic protein feature enrichment in both groups.
